# Estimating the extrinsic incubation period of malaria using a mechanistic model of sporogony

**DOI:** 10.1101/2020.09.30.320044

**Authors:** Isaac J. Stopard, Thomas S. Churcher, Ben Lambert

## Abstract

During sporogony, malaria-causing parasites infect a mosquito, reproduce and migrate to the mosquito salivary glands where they can be transmitted the next time blood-feeding occurs. The time required for sporogony, or extrinsic incubation period (EIP), is a crucial determinant of malaria transmission intensity. The EIP is typically estimated as the time for a given percentile of infected mosquitoes to have salivary gland sporozoites (the infectious parasite life stage). Many mechanisms, however, affect the observed sporozoite prevalence including the human-to-mosquito transmission probability and possibly differences in mosquito mortality according to infection status. To account for these various mechanisms, we present a mechanistic mathematical model (“mSOS”), which explicitly models key processes at the parasite, mosquito and observational scales. Fitting this model to experimental data, we find greater variation in EIP than previously thought: we estimated the range between two percentiles of the distribution, EIP_10_–EIP_90_ (at 27°C), as 4.5 days, compared to 0.9 days using existing methods. This pattern holds over the range of study temperatures included in the dataset. Increasing temperature from 21°C to 34°C decreased the EIP_50_ from 16.1 to 8.8 days and the human-to-mosquito transmission probability from 84% to 42%. Our work highlights the importance of mechanistic modelling of sporogony to (1) improve estimates of malaria transmission under different environmental conditions or disease control programs and (2) evaluate novel interventions that target the mosquito life stages of the parasite.

**Author summary:** *Anopheles* mosquitoes become infected with malaria-causing parasites when blood feeding on an infectious human host. The parasites then process through a number of life stages, which begin in the mosquito gut and end in the salivary glands, where the newly formed infectious parasites can be transmitted to another host the next time a mosquito blood-feeds. The large variability in parasite numbers and development times that exists between mosquitoes, environments and parasites, mean that understanding parasite population dynamics from individual mosquito dissections is difficult. Here, we introduce a mathematical model of the mosquito life stages of parasites that mimics key characteristics of the biology. We show that the model’s parameters can be chosen so that its predictions correspond with experimental observations. In doing so, we estimate key system characteristics that are crucial determinants of malaria transmission intensity. Our work is a step towards a realistic model of within-mosquito parasite dynamics, which is increasingly important given that many recently proposed disease interventions specifically target mosquito life stages of the parasite.

## 1 Introduction

Malaria remains a leading cause of morbidity and mortality worldwide, with an extremely inequitable distribution: over 400,000 people, primarily children under the age of five in sub-Saharan Africa, die annually due to malaria (1). The widespread use of vector control tools that kill adult *Anopheles* mosquitoes is largely responsible for a historical decline in malaria incidence (2); a result foretold by early mathematical models, which predicted the sensitivity of malaria transmission to adult mosquito survival (3,4). For a newly infected mosquito to become infectious, it must survive the extrinsic incubation period (EIP). Since the EIP is long relative to mosquito life expectancy, only older mosquitoes can pass on infection meaning malaria transmission responds acutely to changes in survival (3,5).

The EIP is defined as the duration of sporogony: the obligate reproduction of malaria-causing *Plasmodium* parasites (henceforth parasites) within the mosquito (6). First, female mosquitoes feed on an infectious host. A proportion of these mosquitoes, as determined by the human-to-mosquito transmission probability (5), ingest male and female *Plasmodium* gametocytes within the red blood cells (RBCs) of the blood-meal (7). The change of parasite host (from human to mosquito) involves certain environmental changes, including a decrease in temperature, which collectively trigger gametogenesis and the parasites to emerge from the RBCs (8,9). Fertilisation occurs within the mosquito midgut, where gametes fuse into a single zygote, which differentiates into a motile ookinete (10). Within a few days, ookinetes migrate across the mosquito midgut epithelial wall, and the parasites that survive the mosquito innate immune response (11) go on to form immobile oocysts beneath the midgut basal lamina (12). The number of oocysts remains relatively constant as they grow in size and their genome mitotically replicates (12). Oocysts then burst with each releasing hundreds of infectious sporozoites, which migrate to the mosquito salivary glands, completing the EIP (6,13).

The EIP and human-to-mosquito transmission probability are typically estimated in the laboratory using experimentally introduced infections. Laboratory reared mosquitoes are fed on infectious blood through a membrane feeder, and the parasites are allowed to develop before the mosquitoes are dissected to determine the presence and number of oocysts or sporozoites (13,14). Since a mosquito can be dissected only once, it is not possible to observe parasites dynamics within a single mosquito. Numerous dissections are therefore used to reconstruct the temporal dynamics of sporogony in the population at large (6).

Historically, the EIP has been recorded as the time from blood-feeding until the first mosquito is observed with any salivary gland sporozoites (15,16). Parasite development rates are, however, highly heterogenous due to differences in mosquito nutrition (17,18) and environmental temperature (14,19), and considerable variation remains even after accounting for these factors (14,17). Mosquito and parasite genetic differences may also contribute to variation (15), and even in genetically identical cells reaction rates are noisy, as genes are regulated by small fluctuations in regulatory molecules (20). Given this variation, any single measure – especially one based on a single mosquito, which has historically been the case – does not adequately represent the EIP. Instead, more recent studies attempt to characterise the variability in the EIP, which is reported as the time for given percentiles of infected mosquitoes to display sporozoites (15). A rigorous framework for characterising the distribution of the EIP is yet to be defined.

Not all mosquitoes fed infectious blood will develop observable oocysts or sporozoites. The human-to-mosquito transmission probability depends on many factors including differences in blood meal size, gametocyte density (7,21,22), the mosquito immune response (11,23) and midgut microbiota (24,25) amongst others. To estimate the EIP percentiles, it is necessary to determine the number of infected mosquitoes within the sample as a proportion of all infected individuals. To do this, it is assumed that the maximum observed oocyst- or sporozoite-prevalence is the actual proportion of mosquitoes with viable infection (15). Focussing on raw temporal changes in observed prevalence without an underlying mechanistic model may, however, overlook key processes. In the laboratory, malaria infections may alter mosquito survival in infected mosquitoes (26), meaning the observed maximum prevalence may not represent the true proportion of infected mosquitoes. What’s more, certain variables can impact multiple mechanisms simultaneously (27), requiring a finer-scale understanding of the constituent processes.

Temperature, for example, can modulate parasite development rate (14,19), survival of laboratory mosquitoes (14) and mosquito immune response (28), causing differences in vector competence (29). To date, the relationship between temperature and the EIP has been modelled using the degree-day model, which parameterises the total amount of heat required to complete sporogony (30). Polynomial (quadratic or Brière) functions have also been fit to the relationship between the EIP and temperature (31,32). Parameterisations of both these methods rely on a point estimate of the EIP and do not model the constituent processes that produce these observations. Mechanistic models that simulate parasite population dynamics during sporogony may provide a more informative framework (33,34).

Here, we introduce a mathematical model, the “multiscale Stochastic model Of Sporogony (mSOS)”, to capture the temporal dynamics of *Plasmodium* sporogony. By explicitly modelling the underlying biology, at the parasite, mosquito and experimental scales, we aim to provide an accurate representation of parasite development (35). We fit mSOS to laboratory membrane feeding assay data and present new EIP estimates which contrast with the existing literature. We also demonstrate how mSOS can be used to estimate the impact of temperature on *Plasmodium falciparum* sporogony within *Anopheles* mosquitoes. All code and data needed to recapitulate our results is available at https://github.com/IsaacStopard/mSOS.

## 2 Models and methods

### 2.1 Data

We are unaware of a single study with sufficient data to fit the model. To capture the full population dynamics of oocysts and sporozoites, we combined data from four published studies (Table 1). Most data came from Standard Membrane Feeding Assays (SMFA) from the same laboratory, conducted with the same parasite-vector combination (14,18,36). These studies collected data over a range of days and temperatures using a laboratory strain of *P. falciparum* (NF54) and laboratory-reared *Anopheles stephensi* mosquitoes. To examine oocyst prevalence prior to day five, we also included data from a Direct Membrane Feeding Assay (DMFA) (37), which used wild *P. falciparum* parasites to infect laboratory-reared *Anopheles gambiae sensu stricto* mosquitoes. It is assumed there is no difference between the two parasite-vector combinations. All studies replicated the following experimental protocol: mosquitoes were raised from larvae under laboratory conditions; adult female mosquitoes were then fed on *P. falciparum-*infected blood via a membrane feeder; at various days post infection, blood-fed mosquitoes were collected, dissected and the presence of oocysts or salivary-gland sporozoites identified by microscopy. The oocyst load within each mosquito was estimated by counting observable oocysts. Sporozoites were quantified by presence or absence. Only mosquito larvae kept on a high food diet were used in these analyses, and, in all cases, mosquitoes were housed at a constant temperature and humidity (see Table 1 for the range of different temperatures explored).

**Table 1.** Summary of different studies, parasite-vector combinations and experimental proceedures used to parameterise the model.

| Malaria species (strain) | Vector species (strain) | Days post blood-feeding when oocyst presence or load was determined by dissection | Days post blood-feeding when sporozoite presence was determined by dissection | Dead mosquitoes counted? | Adult mosquito housing temperature (°C) | Reference |
| --- | --- | --- | --- | --- | --- | --- |
| *P. falciparum* (NF54) | *A. stephensi* (laboratory) | 7 (oocyst load) | 15 | Yes, infectious blood-fed & uninfected blood-fed (control) mosquitoes | 27, 30 & 33 | Murdock et al. (36) |
| *P. falciparum* (NF54) | *A. stephensi* (laboratory) | Not recorded | Varied with temperature: 10 to 23 & 25 (21°C), 9 to 23 & 25 (24°C), 8 to 18, 20, 22, 24 & 25 (27°C), 6 to 17 & 19 (30°C), 5 to 15, 17 & 19 (32°C) and 5 to 14 & 16 (34°C). | Yes, infectious blood-fed mosquitoes only | 21, 24, 27, 30, 32 & 34 | Shapiro et al. (14) |
| *P. falciparum* (NF54) | *A. stephensi* (laboratory) | 5, 6, 7, 8, 9 & 10 (oocyst load) | 9, 10, 11, 12, 13, 14, 15 & 16 | Yes, infectious blood-fed mosquitoes only | 27 | Shapiro et al. (18) |
| *P. falciparum* (wild) | *A. gambiae* s.s. (laboratory) | 3, 4, 5, 6, 7 & 8 (oocyst presence used) | Not used | No | 27 - 28 | Bompard et al. (37) |

### 2.2 Model overview

mSOS models the oocyst and salivary gland sporozoite (henceforth sporozoite) life stages, as oocysts are the most widely recorded outcome of membrane-feeding assays, and sporozoites are the most epidemiologically important life stage. We explicitly model the counts of oocyst- and sporozoite-positive mosquitoes to determine how the prevalence (i.e. the percent of positive mosquitoes among the sample) of each life stage varies over time. We also model oocyst intensity (mean oocyst load among the sample), as it provides more information on the underlying parasite dynamics and is considered a more reliable estimate of human-to-mosquito transmission (38,39).

Sporogony is modelled at multiple scales, and all parameters are fit simultaneously in a single model. At the parasite scale, we model the sequence of three life stages: inoculant → oocyst → sporozoite (Figure 1A). We do not model the pre-oocyst life stages since they are harder to count and are rarely recorded. Rather, these are represented in a notional “inoculant” stage ingested during blood-feeding, each of which develops into a single oocyst. mSOS captures both the time it takes for each oocyst to appear, and the time for each oocyst to develop into sporozoites. The time taken for each of these transitions to occur is assumed independent of the other, and we model these times stochastically.

**Figure 1.**
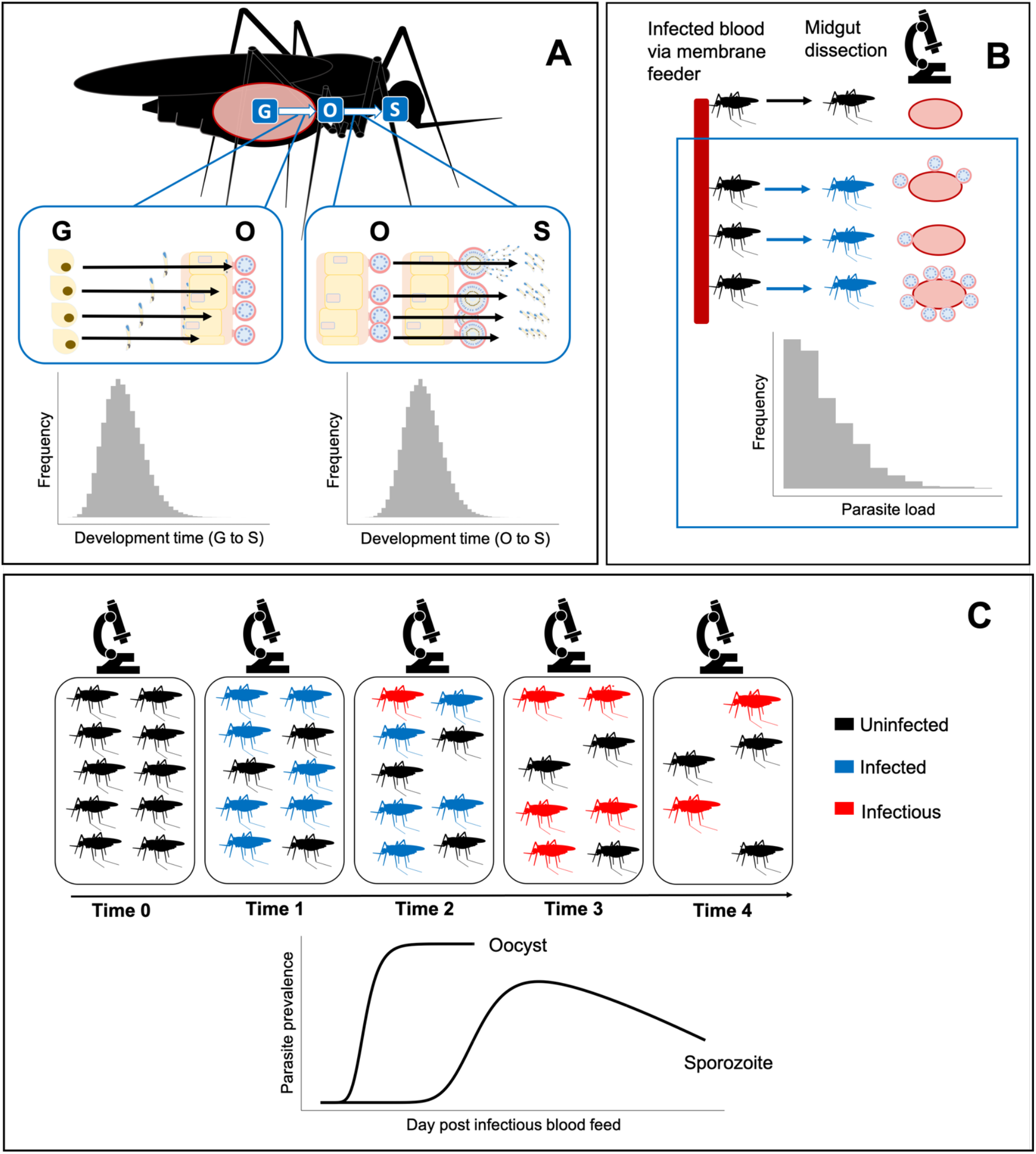
Structure of mSOS: a multiscale model of the population dynamics of *Plasmodium falciparum* during sporogony. (a) Multiple malaria parasites are found within a single mosquito; we separately model development time from inoculation at blood-feeding (G) to oocyst (O) and from oocyst to salivary-gland sporozoites (S). If dissected, a mosquito is “positive” for a particular parasite life stage if at least a single parasite has developed. We do not model the decline in observed oocyst numbers due to oocyst bursting, since we do not have sufficient later oocyst observations. (b) Mosquitoes are infected via a membrane feeder; parasite load varies in each due to differences in the number of parasites ingested and variation in mosquito immune response. (c) Temporal dynamics of sporozoite prevalence within a mosquito population: following the infectious blood-feed, a proportion of the population is infected with malaria parasites. The parasites develop into sporozoites causing the mosquitoes to become infectious. Throughout the experiments, mosquito mortality (in the laboratory) may be greater in infected mosquitoes, resulting in an eventual decline in observed sporozoite prevalence.

At the mosquito scale, mosquitoes are initially infected with different numbers of inoculant stage parasites, with each load determined stochastically. We also allow infectious blood-fed mosquitoes to avoid infection (via a human-to-mosquito transmission probability) (Figure 1B). Potential differences in survival of the mosquito population due to differences in malaria infection (26) are explicitly modelled.

At the observation level, to reconstruct the temporal dynamics of sporogony at a population level, we recreate populations of parasites nested within a population of mosquito vectors, which then form samples of dissected mosquitoes (Figure 1C). We allow the composition of each sample of dissected mosquitoes to vary according to the proportions of infected and uninfected mosquitoes alive at the time of dissection.

To incorporate temperature-dependency, we allow the time for the appearance of oocysts (40,41), and the human-to-mosquito transmission probability, to be influenced by temperature. The development time from oocyst to sporozoites and the mean parasite load among infected mosquitoes are assumed to be independent of temperature.

In what follows, we describe the model in detail. To provide greater clarity, a Mathematica file that steps through the derivations is available in the supplementary information.

#### 2.2.1 Parasite scale

We model the time taken, *T_GO_*, for each undeveloped oocyst at the time of blood-feeding (the notional inoculant stage denoted ‘*G*’) to develop into an observable oocyst (‘*O*’). We then model the time taken, *T_OS_*, for each oocyst to develop, burst, and for the sporozoites to migrate to the salivary glands (we term sporozoites in the salivary glands ‘*S*’). So, these transitions follow:

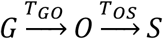

These development times are modelled stochastically as: 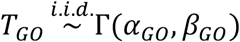 and 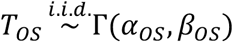; that is, we assume that the time it takes a subsequent oocyst to burst into sporozoites is independent of the initial time taken for the oocyst to develop. This assumption also implies the transitions of each parasite occur independently, meaning density dependent population dynamics do not occur. We assume a parameterisation of the gamma distribution such that its mean is *E*(*T_i_*) = *α_i_*/*β_i_*, where *i* ∈ (*GO,OS*), *α_i_* is the shape parameter and *β_i_* is the rate parameter.

The time required for *G* to develop into *S* is given by the sum: *T_GS_* = *T_GO_* + *T_OS_*. An analytic form for the cumulative density function (CDF) of the sum of two gamma distributed random variables with different rate (*β*) parameters is currently not known. This aggregate distribution can, however, be approximated by a gamma distribution where the mean, *λ* and variance, *σ^2^* match that of the true distribution (42),

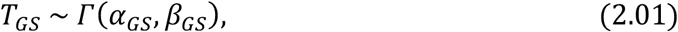

where,

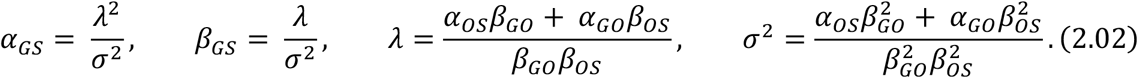

#### 2.2.2 Mosquito scale

Mosquitoes are treated as self-contained populations of parasites. When feeding on infectious blood, mosquitoes receive a heterogeneous load of gametocytes, with some receiving none at all (7). Additionally, not all parasites develop into the observed oocyst stage due to the innate immune response of some mosquitoes (13,38). Evidence indicates that, assuming the mosquito is still alive, the majority of oocysts will produce sporozoites (43). Hence, we term a mosquito “successfully infected” if at least one viable oocyst could develop from the initial infective load given sufficient time and define this probability of successful infection (i.e. the human-to-mosquito transmission probability) as *δ*. Among successfully infected mosquitoes, the initial load of G-stage parasites, *n*, is modelled by a zero-truncated negative binomial distribution,

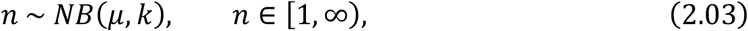

where, *μ* is the mean and *k* is the overdispersion parameter. Evidence indicates *μ* and *δ* may be jointly influenced by gametocyte density, which is modulated by temperature (29). We treated these parameters as independent, as there was an insufficient combination of gametocyte density and temperature treatments in the data.

To model the observed oocyst or sporozoite presence, we assume a mosquito is measured as “positive” if at least a single parasite has developed to the given life stage. That is, we assume that dissection always uncovers some parasites of a given life stage should any exist.

Let the time taken for parasite *j* to develop into a subsequent stage be *T_j_* for *j* = 1,2, …, *n* parasites within a specific mosquito. Whether a mosquito is positive for a particular stage at time *t* then depends on whether 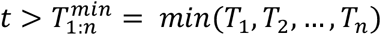. Since, we are concerned with the minimum of a series of random variables, we are in the realm of order statistics – see (44). Specifically, 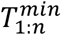 is the 1^st^ order statistic of the sample of *n* development times. Considering a single parasite, *j*, the probability that it has developed by time *t* is given by *Pr*(*t*≥ *T_j_*) = *Q*(*α*, 0, *βt*), where *Q*(*α*, 0, *βt*) is the CDF of the gamma distribution governing that particular life stage transition. Specifically, *Q* is the generalized regularized incomplete gamma function 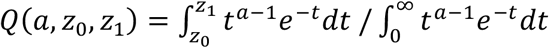, and *α, β* are the shape and rate parameters of the underlying gamma distribution (these parameters will, in general, be different for *T_GO_* and *T_GS_*). Considering *n* parasites within a single mosquito, the probability that at least one of them has developed by time *t* is:

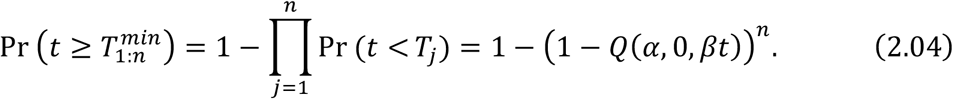

To model the number of oocysts at a given time within an individual infected mosquito, *Y*(*t*), we track the number of G-stage parasites, from the initial load *n*, that have developed by a given time, which follows a binomial distribution,

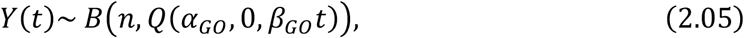

where *Q*(*α_GO_*, 0, *β_GO_t*) is the cumulative probability that an individual parasite has developed into an oocyst by time *t*.

Mosquitoes die throughout the course of the experiments, which we explicitly model via a Cox regression model. We estimate the probability a mosquito is alive immediately prior to time *t*, *A*(*t*), by fitting this model to the mosquito survival data (Table 1) and allow for differences in the survival due to infection by comparing groups of mosquitoes exposed to uninfected and infected blood and allowing the hazard to vary with infection status. Mosquitoes exposed to infected blood may not actually become infected, so this group consists of a mix of infected and uninfected individuals. By fitting the complete model, we account for this to estimate infection-specific differences in mortality, see S7.1.1 for full details. A previous analysis of a subset of the data determined that the Gompertz distribution, in which the mosquito mortality rate increases with age (senescence) (14), was the best fit, and we use this distribution here.

#### 2.2.3 Observation level

During the experiments, samples of mosquitoes are dissected at particular time intervals. From these samples, two possible measurements are recorded: (1) the aggregate count of parasite-positive mosquitoes (either oocysts or sporozoites) and (2) the number of oocysts counted in each of the dissected mosquitoes.

We model the aggregate count of positive mosquito data thus. At each time, a sample of mosquitoes, *D*(*t*), are dissected, likely comprising a mix of infected and uninfected individuals. The number of infected mosquitoes within the sample, *I*(*t*), is modelled as a binomial random variable,

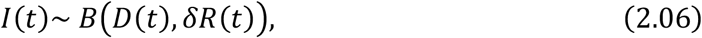

where *δ* is the probability successful infection occurs during the infectious blood-feeding, and *R*(*t*) is the fraction of infected mosquitoes alive. *R*(*t*) is the ratio of survival probabilities for infected (*E*) and uninfected (*U*) mosquitoes:

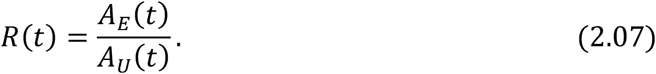

Note, we assume that once a mosquito is infected it will remain so for the duration of its lifespan (45).

A mosquito may be successfully infected but no parasites observed during dissection, if insufficient time has passed since blood-feeding. If *n* was known for each dissected mosquito, the probability dissection would detect parasites of a given life stage, at time *t*, is given by eq. (2.04). In reality, *n* is not known. We model this uncertainty using eq. (2.03) and incorporate it into the probability any infected mosquito has observable parasites by marginalising *n* out of the joint distribution,

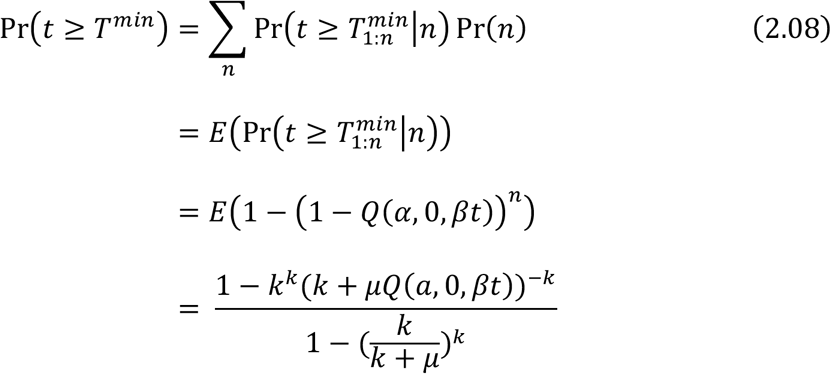

where *T^min^* is the time when first parasite of the given life stage appears (*O* or *S*), and *E_n_*(.) denotes the expectation with respect to the distribution of *n* (eq. (2.03)).

The count of infected mosquitoes in which the parasite life stage of interest (*O* or *S*) is observed, *X*(*t*), given a sample of infected mosquitoes, *I*(*t*), is then also binomially distributed,

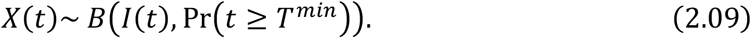

Within a given sample of mosquitoes, the number of successfully infected mosquitoes, *I*(*t*), is not known – we only observe the count of observed infected individuals, *X*(*t*), and the total number dissected, *D*(*t*). We incorporate this uncertainty in *I*(*t*) by marginalising this quantity out of the joint distribution defined in eqs. (2.06) and (2.09), resulting in the following binomial distribution describing the observed count,

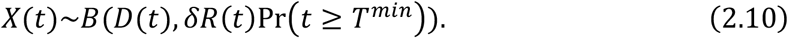

We next detail how our model describes oocyst counts, *Y*(*t*). A particular mosquito may yield a zero count when dissected either if it was not successfully infected or if insufficient time has elapsed for any oocysts to develop. The probability a sampled individual is successfully infected is *δR*(*t*). The count of oocysts to have developed by a given time within a successfully infected mosquito is described by eq. (2.05): this probability distribution depends on *n* (the number of G-stage parasites) – an unknown quantity. To derive the sampling distribution of oocyst counts, we marginalise *n* out of the joint distribution assuming its uncertainty is described by eq. (2.03). Combining this with our underlying uncertainty regarding the mosquito’s infection status results in the following expression describing the probability of counting *Y* oocysts at time *t*, in a dissected mosquito,

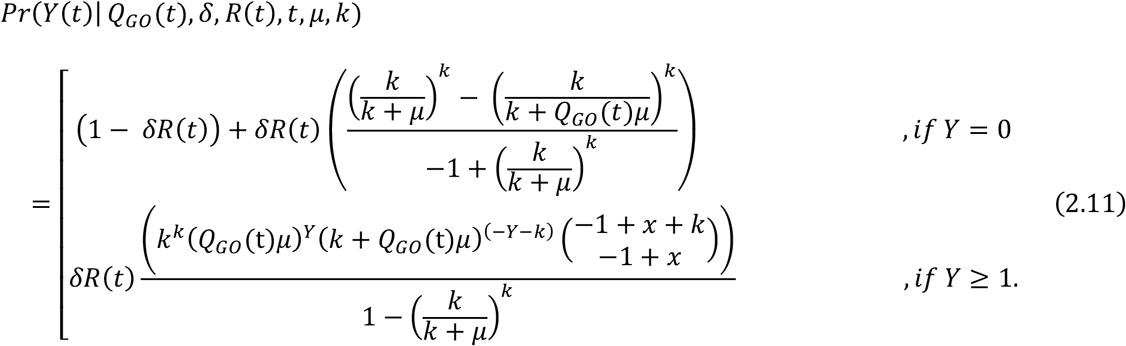

where, for notational convenience, we term *Q_GO_*(*t*) = *Q*(*α_GO_*, 0, *β_GO_t*). From eq. (2.11), the mean oocyst intensity within the population is given by 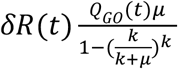. Mosquitoes are typically dissected for oocysts before any burst, so we do not model how bursting would lead to a decline in oocyst prevalence or intensity.

### 2.3 Incorporating temperature-dependence

First, we fit the model to data collected under standard insectary conditions (27 °C), which included the complete range of oocyst intensity, oocyst prevalence, sporozoite prevalence and survival (grouped by infectious blood-fed and control) data. Next, to investigate the impact of temperature on the EIP, we fit the model to the data collected at each other temperature, which included sporozoite prevalence and survival data only. Due to the lack of oocyst data, development times from day of blood-feeding to sporozoite were estimated as a single gamma distribution with shape, *α_GS_*, and rate, *β_GS_*. Data collected at 33°C was excluded as mosquitoes were only sampled on a single day. We refer to these individual temperature model fits collectively as the “single temperature models”.

To estimate a functional relationship between temperature and EIP, we fit the model to all the data simultaneously (the “all temperature model”). To do so, we first plotted the *α_GS_* and *β_GS_* parameters of the single temperature models against temperature (Figure S1). This indicated an approximate linear relationship between temperature and the development rate, *β_GS_*. Laboratory studies have found the early part of sporogony of laboratory strains of *P. falciparum*, in *A. stephensi*, is most temperature sensitive (40,41), so we assumed the temperature variation affected the rate at which parasites develop from *G* to *O*. The development rate was then modelled as a linear function of temperature, *C*,

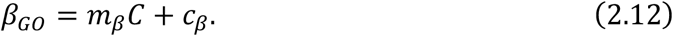

The infection of mosquitoes has been found to decline at high temperatures, in the laboratory, due to factors such as the impact of temperature on vector immunity (27) and the establishment of gametocytes (29). To account for this, we included a relationship between temperature and the human-to-mosquito transmission probability, δ, modelled as a logit-linear function,

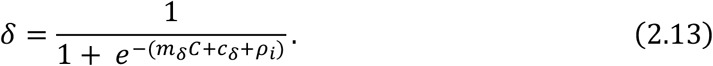

To capture differences between membrane feeding assays (38), we included a hierarchical term *ρ_i_*, in eq. (2.13) where,

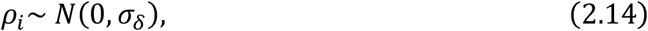

and *i* refers to an individual experiment, defined as any unique combination of study and temperature treatments (i.e. the combination of reference and adult mosquito housing temperature values in Table 1).

Kaplan-Meier plots of mosquito survivorship and existing studies indicated that laboratory *A. stephensi* survival depended on temperature (Figure S2). To account for this, a temperature-dependent mortality term was incorporated into the Cox model hazard,

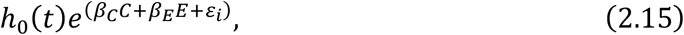

where C is temperature, E is the infection status (*E* = 1 indicates successful infection; and *E* = 0 for uninfected individuals), and *h*_0_(*t*) is the baseline hazard modelled by a Gompertz distribution. To account for experimental heterogeneity in mosquito survival between experiments, a hierarchical error term, *ε_i_*, was added to eq. (2.15) where,

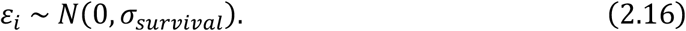

To facilitate model fitting, we scaled the temperature values such that temperature was centred around 0 with a standard deviation of 1.

### 2.4 Model fitting

The models were fit under a Bayesian framework, using the probabilistic programming language Stan (46), which implements the No-U-turn Markov chain Monte Carlo (MCMC) sampler (47). We specified weakly informative priors following a literature search (Table S1). For each model fit, four chains were run, with 1500 warmup iterations and 4500 iterations in total. Convergence was determined by 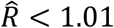 for all parameters, and bulk and tail effective sample sizes (ESS; an estimate of sampling efficiency) greater than 400 (see Table S2 for these values). We visually examined the trace plots (Figure S3) and illustrate the difference between prior and posterior distributions in Figure S4.

### 2.5 Model investigation

To compare our estimates with the predominant approach, we estimated the EIP and human-to-mosquito transmission probability using a logistic model (14,15,18,27,48), which we fit using a non-linear least squares approach (see supplementary information 7.1.3). Given a decline in the observed sporozoite prevalence at later time points, the data was subset to include only data points before the time of peak observed sporozoite prevalence, as has been done previously (14).

Data did not exist to investigate the impact of parasite load experimentally. We therefore used our fitted model to simulate the temporal dynamics of sporogony with different mean parasite loads (among infected mosquitoes), holding all other parameters at their mean posterior values for 27°C (sensitivity analysis).

## 3 Results

### 3.1 Temporal dynamics of sporogony

We first considered the observed temporal dynamics of sporogony at a single temperature (27°C; standard insectary conditions). All values we report are the posterior predictive means and the credible intervals are the 95% central posterior estimates; times reported are the number of days post infectious blood-feed. From the model fit, we estimate that oocyst prevalence was 10% at 2.2 days (CI: 2.1–2.3 days) and peaked at 69% (CI: 68–71%) after approximately 5.9 days (CI: 5.8–6.0 days) (Figure 2a). The modelled oocyst intensity peaked at 2.0 (CI: 1.9–2.1) oocysts per mosquito, shortly after oocyst prevalence peaked (Figure 2b). In the raw data, sporozoite-positive mosquitoes were first observed on day 10 when the sporozoite prevalence was 1.4%; our model estimates are slightly earlier: the modelled sporozoite prevalence reached 10% at 9.5 days (CI: 9.3–9.7 days) (Figure 2c). The time required for sporozoites to appear varied between mosquitoes, and, only after 15.9 days (CI: 14.2–16.0 days) did the sporozoite prevalence peak at 63% (CI: 61–65%).

**Figure 2.**
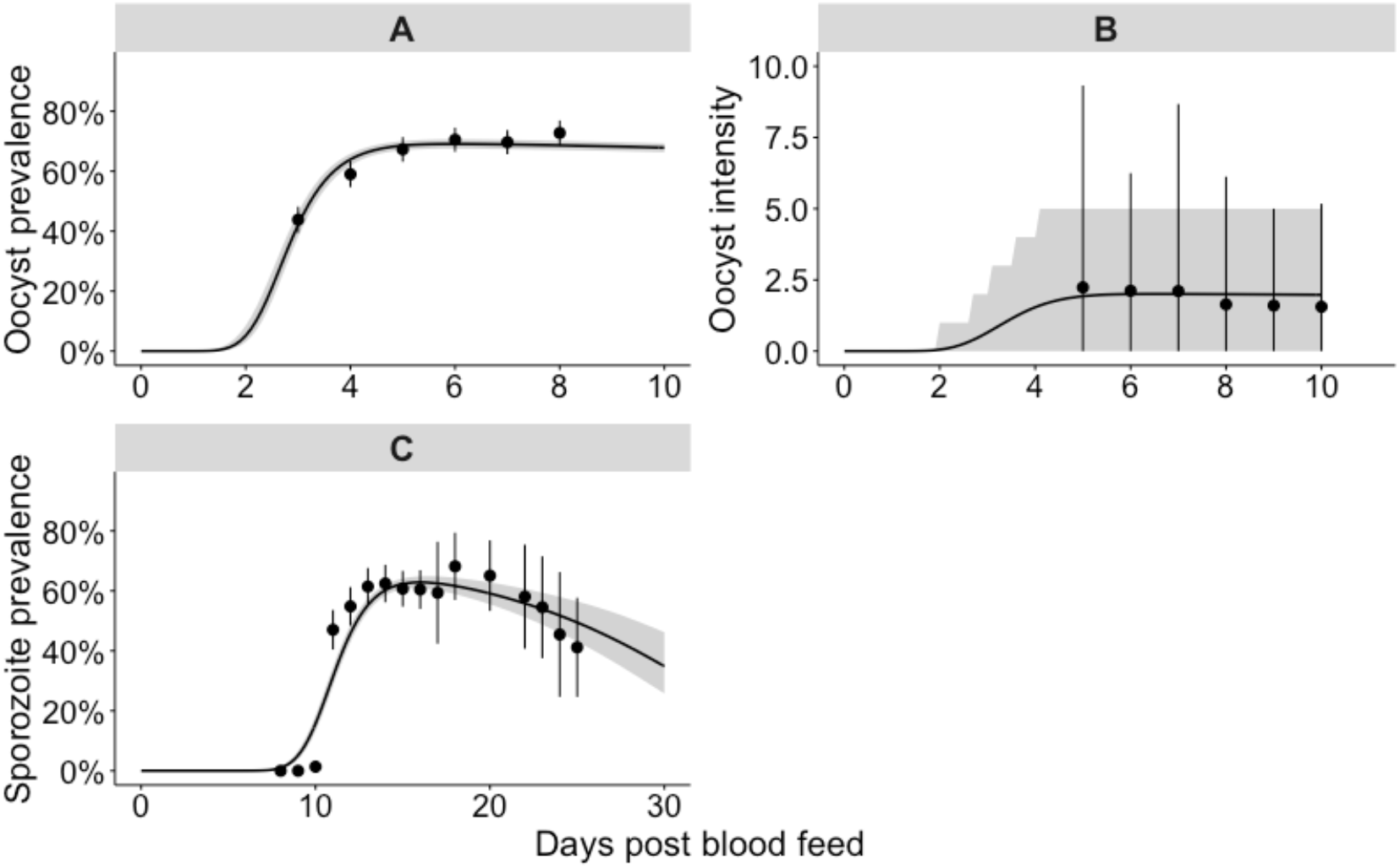
Single temperature 27°C model fit to the oocyst and sporozoite data. The panels show our model fit to the 27°C dataset: panel A to the oocyst prevalence, panel B to the oocyst intensity data and panel C to the sporozoite prevalence. (A & C) Points: parasite prevalence of the laboratory mosquito data (95% confidence intervals are given by the point range). The grey shaded area represents the 95% credible interval of the model posterior predictive means, the median posterior predictive mean is shown by the black line. (B) The points show the mean parasite load among all blood fed mosquitoes (intensity), the point range indicates the 2.5%–97.5% quantiles of the raw data. The shaded area represents the 2.5%–97.5%-quantiles of the negative binomial distribution; where the location and overdispersion parameters at set to their posterior means.

At the individual parasite level, we consider the time taken for each modelled parasite life stage transition to occur: the mean time taken was 3.4 days (CI: 3.3–3.6 days) for G to O, 9.2 days (CI: 8.9–9.5 days) for O to S and 12.6 days (CI: 12.3–12.9 days) for G to S (Figure S5a). The majority of individual parasites (99.5%) had undergone each of these transitions at 6.1 days (CI: 5.6–6.6 days) for G to O; 14.7 days (CI: 13.9–15.5 days) for O to S; and 18.5 days (CI: 17.8–19.3 days) for G to S (Figure S5b).

### 3.2 Impact of temperature

We next considered the impact of temperature on sporogony. The *single temperature models* fitted the sporozoite prevalence data well across the entire range of temperatures (Figure S6). *The all temperature model* fitted the sporozoite prevalence data well at lower temperatures (between 21°C and 30°C); at higher temperatures, the model tended to overstate the EIP (Figure 3). The survival of both infectious blood-fed and control mosquitoes was only available at 27°C, 30°C and 33°C, and consequently the *single temperature model* survival parameters had greater freedom to vary. Indeed, the *all temperature model* (Figure S7) fits to the survival data were less variable and more predictable across different temperatures than the *single temperature models* (Figure S8). Consequently, here we provide the *all temperature model* (Figure 3, Figure S9) results due to their greater generalisability.

**Figure 3.**
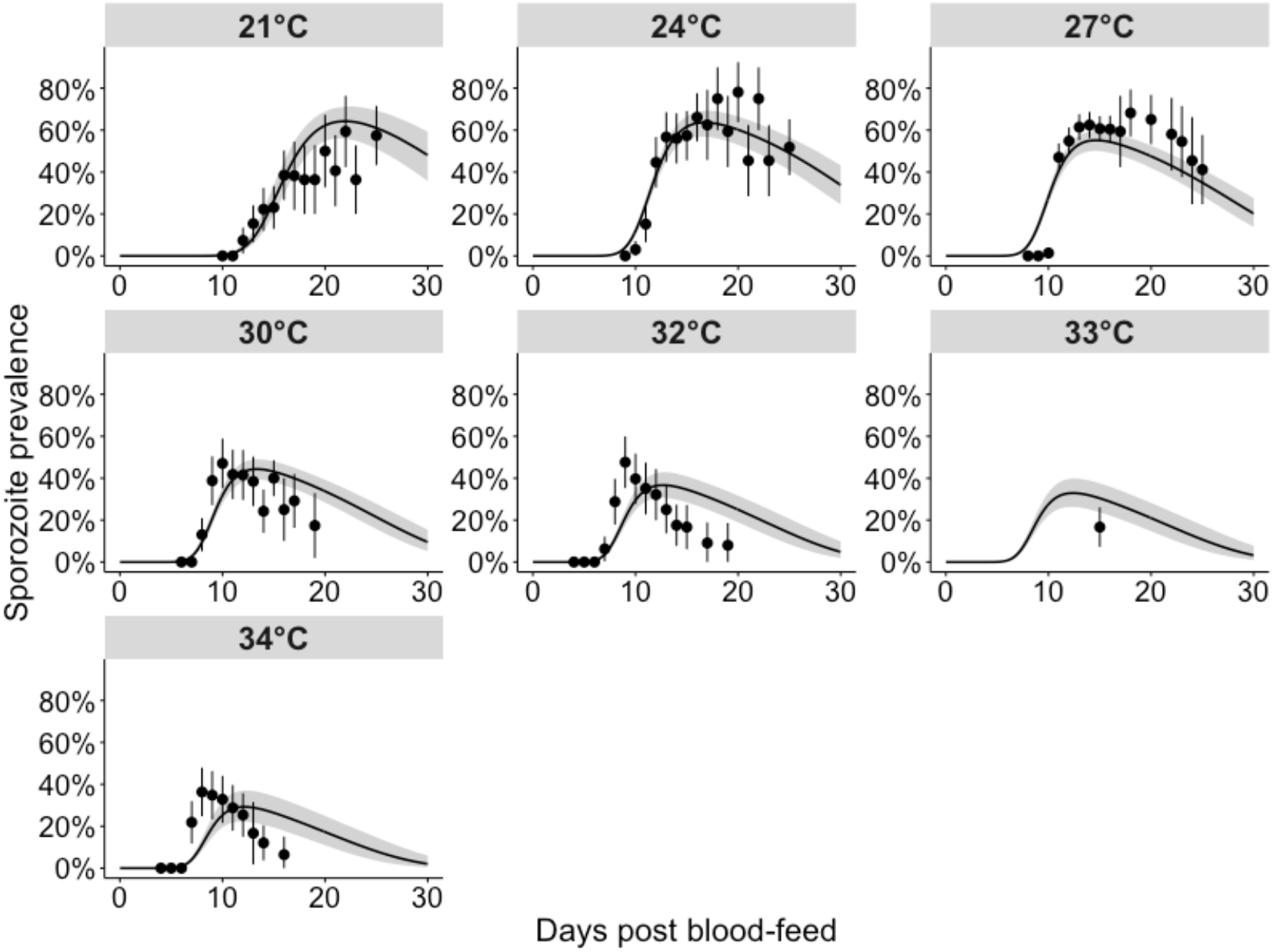
Model fits to sporozoite prevalence data across all temperatures. These fits were generated by fitting a single model to all temperatures simultaneously (“all temperature model”), with the functional form of temperature as described in eqs. (2.11) and (2.12). Black points: parasite prevalence of the laboratory mosquito data (95% confidence intervals are given by the vertical black lines). The grey shaded area represents the 95% quantiles of the posterior predictive means; the black lines represent the median posterior predictive means.

For the *all temperature model*, infection resulted in a higher risk of mosquito death, with a hazard odds ratio of 1.65 (CI: 1.47–1.86). Increases in temperature elevated mosquito mortality with a hazard odds ratio of 1.57 (CI: 1.33-1.86) per 3.5°C change. At 10 days post-infection, the probability of an infected mosquito being alive, *A(t)*, was 0.91 (CI: 0.87–0.94) at 21°C versus 0.67 (CI: 0.58–0.75) at 34°C, with similar differences for uninfected individuals.

At higher temperatures, fewer mosquitoes develop sporozoites, but those that do, do so faster. We estimate that, between 21°C and 34°C, increases in temperature reduced the EIP: EIP_10_ fell from 12.9 days (CI: 12.5–13.3 days) to 6.9 days (CI: 6.7–7.1 days), EIP_50_ declined from 16.1 days (CI: 15.5–16.8) to 8.8 days (CI: 8.6–8.9 days) and EIP_90_ fell from 20.4 days (CI: 19.4–21.8 days) to 11.3 days (CI: 11.0–11.7 days) (Figure 4a). Table S3 provides the EIP percentiles at all temperatures. Increases in temperature reduced the human-to-mosquito transmission probability, which fell from 84% (CI: 76–89%) at 21°C to 42% (CI: 34– 50%) at 34°C (Figure 4).

**Figure 4.**
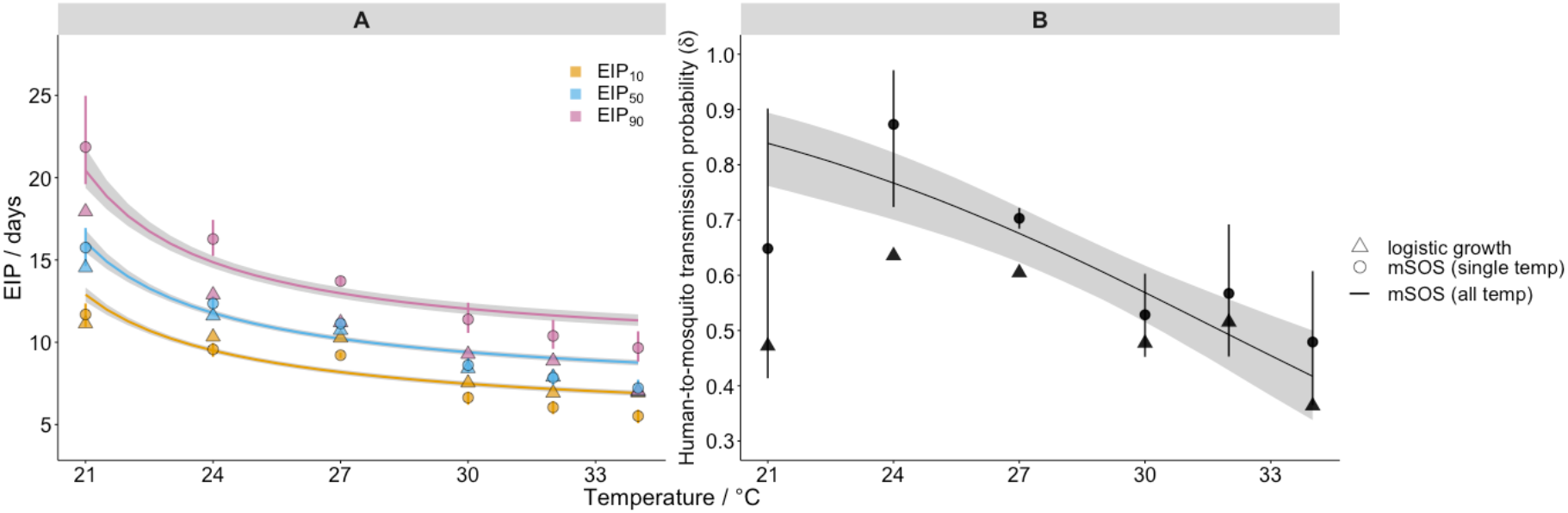
Effect of temperature on malaria transmission parameters. Panel A shows the model impact of temperature on the EIP quantiles (as indicated in legend); Panel B shows its impact on the human-to-mosquito transmission probability. In both panels, the lines show impact as estimated by the *all temperature* mSOS model with 95% posterior intervals indicated by shading; the discrete round points show independent estimates of the *single temperature* mSOS model at each temperature, 95% posterior intervals are shown by the vertical lines. The number of iterations used to calculate the EIP plot were thinned to every 5^th^ iteration for efficiency. The discrete triangle points show the logistic growth model parameter estimates at each temperature.

### 3.3 Key differences in transmission parameters estimated using mSOS

Next, we compared our estimates of two important malaria transmission parameters – the EIP and human-to-mosquito transmission probability – to those obtained using the predominant literature approach based on logistic regression (logistic model fits are shown Figure S10). In our framework, we can disentangle the development time of parasites from other underlying processes, specifically mosquito mortality induced by malaria infection, which can influence the observed sporozoite prevalence. In doing so, we quantify the EIP distribution as the time taken for a given percentile of the infected mosquitoes to display sporozoites, in the absence of other mechanisms that determine the observed sporozoite prevalence. This is given by the time at which the cumulative probability an infected mosquito develops any salivary gland sporozoites, Pr(*T*_1_ ≤ *t*), reaches a given value. Using this approach, across all temperatures, the variation in the EIP distribution is greater than the equivalent logistic model estimates (Figure 4a). At 27°C, for example, our *single temperature* model estimated an EIP_10_–EIP_90_ range of 9.2–13.7 days compared to 10.3–11.2 days estimated by the logistic approach.

The mSOS estimates of the human-to-mosquito transmission probability were consistently higher than the equivalent values estimated by the logistic model (Figure 4b). The 27°C *single temperature* model estimate, for example, was 70% (CI: 68–72%) as opposed to the logistic growth model estimate of 60%. This is because, in the laboratory data we analysed, infected mosquitoes died at an elevated rate, meaning that the proportion of infected mosquitoes declined with time and the observed peak of sporozoite prevalence is not representative of the initial proportion of infected mosquitoes. Not accounting for these mechanisms results in an underestimate of the human-to-mosquito transmission probability.

### 3.4 Sensitivity analysis of the mean parasite load

Within our model, the time-taken for each simulated parasite to develop is both independent and stochastic. At each time point, whether or not that parasite has yet developed can be viewed as a toss of a coin: the more coins are tossed, the more likely it is that one will land on heads, or equivalently a parasite will have developed, by chance. Higher parasite loads then lead to earlier rises in parasite prevalence. Figure 5A shows the simulated sporozoite prevalence over time across populations with different mean parasite load parameter values, which indicates within the model greater parasite numbers cause the prevalence to peak earlier. Figure 5B shows the resultant EIP quantiles as a function of parasite load: increasing the mean parasite load from 1.7 to 25.0 reduced the modelled EIP_50_ from 10.9 to 8.0 days.

**Figure 5.**
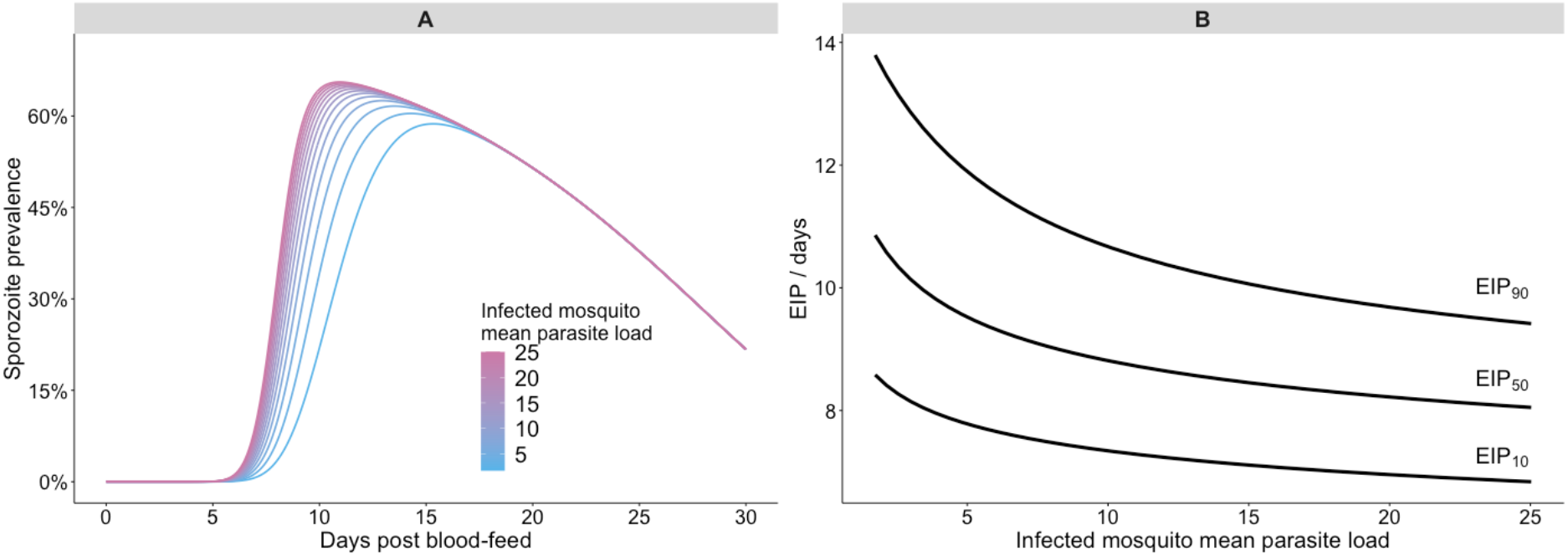
Modelled impact of parasite load on the extrinsic incubation period. Panel A shows the impact of varying the mean parasite load of infected mosquitoes on the temporal dynamics of sporozoite prevalence in a sensitivity analysis; panel B summarises how the EIP is affected by the same parameter in the sensitivity analysis. All other parameters were held constant at their mean posterior values.

## 4 Discussion

Membrane feedings assays form the bulk of experiments used to determine key parasitological parameters of malaria transmission, but despite the seeming simplicity of these experiments, numerous hidden processes contribute to the observed data. Here, we applied a systems biology approach to develop a multiscale model of the temporal dynamics of sporogony (mSOS) that explicitly account for these underlying processes. To illustrate its use, we fitted mSOS to experimental data and demonstrated its predictive accuracy across a range of experimental protocols. In doing so, we estimated two important determinants of malaria transmission intensity – the EIP and human-to-mosquito transmission probability. By adopting a mechanistic approach, our EIP estimates are defined in terms of the underlying biology rather than characteristics of the raw experimental data. Our estimates indicate greater variation in the EIP than previously thought and highlight the importance of accounting for this variation when making epidemiological predictions (see also (15)). This variation could be included by embedding mSOS within transmission dynamics models of malaria.

The influence of parasite-induced mosquito mortality is still a source of debate, and likely to vary depending on the parasite-vector system (26,36). Nonetheless, accounting for potential differences in mortality is essential to avoid bias in estimates of both EIP and the human-to-mosquito transmission probability. By modelling mosquito infection and survival in a single framework we could estimate malaria infection induced differences in mosquito survival, whilst accounting for mosquitoes that do not develop oocysts within the infectious blood fed group. At higher temperatures, we found that these differences only partly explained the sporozoite prevalence data. Another study observed no difference in the survivorship of wild caught *A. gambiae* mosquitoes due to malaria infection, yet *P. falciparum* detection did decline over time (49). So, it is possible that mosquito survivorship was not behind these observed declines and alternative explanations, such as sporozoite loss during sugar-feeding (50,51) or sporozoite mortality could require further investigation.

Temperature is an important predictor of *P. falciparum* prevalence among humans, its impact is, however, often non-linear and location-specific (52–54). Differences in the EIP due to temperature, parameterised using laboratory data, are commonly used in the prediction of the spatiotemporal limits and endemicity of malaria transmission (32,55,56). Over the temperature range investigated, our estimates qualitatively match those of (15), which determined that the EIP falls with increases in temperature, and that the incremental change is greatest at lower temperatures. In a semi-field setting, increases in temperature, due to the interaction between deforestation and altitude, similarly decreased the EIP of wild *P. falciparum* (19), but very little field data exists. To more accurately simulate the EIP in the field will require consideration of the impact of the time of biting (57), diurnal temperature fluctuations (58) and mosquito resting location on the temperatures parasites are exposed to during different life stages. The degree to which the EIP of locally adapted strains differ to the laboratory strains must also be considered (59). Current estimates indicate climate change has the potential to change the number of people at risk of malaria (56,60,61), though there is uncertainty due to the complexity of how the parasite and mosquito will interact with the local changing environment in the long term. Characterising spatiotemporal heterogeneity in the EIP in the field is nonetheless critical to assess our current understanding of malaria and its control as well as the changing risk of malaria resulting from the interactions between climatic, land-use (62) and socio-economic factors (63).

Within our model, heterogeneity in the EIP emerges from differences in the development times of individual parasites. Density-dependent processes (64,65) and intraspecific variation in individual mosquito characteristics, such as body size (66), may also impact the observed sporozoite prevalence. To include such processes in the model in a data-driven way requires higher resolution data to differentiate the impact of different hypotheses. To parameterise mSOS, we collated data from four previously published experimental infections. We therefore recommend that future studies dissect mosquitoes for both oocysts and sporozoites across a greater range of times post infection. It would also be useful if mosquito mortality were recorded, ideally determining the past infection status of carcases through molecular methods.

The work shows how higher parasite loads within a mosquito might decrease the EIP. To our knowledge, this possible relationship between parasite load and development time has not been considered elsewhere but could have potentially important epidemiological consequences as it indicates that onwards transmission may be more efficient from more infectious people. Further work is needed to explore this hypothesis given that our model assumes the EIP is determined on a per-oocyst basis and that developmental times are independent within an individual mosquito which needs to be verified experimentally. What’s more, other processes may be operating: emerging experimental evidence hints that a decrease in resource availability per parasite (at high parasite loads), as determined by the parasite load and the number of times the mosquito blood-feeds, may decrease the parasite development rate (48,67). Since (a) transmission is highly sensitive to changes in EIP, (b) transmission blocking interventions cause a decline in oocyst intensity (38), and (c) parasite load in the field may be higher than previously thought (37), the influence of parasite load on EIP merits further investigation.

It is now well over a century since *Plasmodium* parasites were first uncovered in dissected anopheline mosquitoes. Today, we remain dependent on dissection for understanding the parasite lifecycle in mosquitoes, and there remains much to learn. The model we introduce here provides a new way to parse dissection data to probe the underlying biology. Our model can be extended to incorporate more fine scale characteristics of parasite ecology, but to do so in a principled manner requires more fine scale data. As such, we foresee a great necessity and opportunity for closer collaboration between experimentalists and modellers in the future.

## 5 Acknowledgements

We would like to thank Professor Matthew Thomas for useful insight and the Thomas lab at Penn State University for producing and publishing the high-quality data upon which our study is based.

IJS would like to thank the Natural Environment Research Council (NERC) for PhD funding administered through the Quantitative Methods in Ecology and Evolution (QMEE) Centre for Doctoral Training (grant reference: NE/P012345/1). The authors would like to acknowledge the MRC Centre for Global Infectious Disease Analysis (grant reference: MR/R015600/1), this award is jointly funded by the UK Medical Research Council (MRC) and the UK Department for International Development (DFID) under the MRC/DFID Concordat agreement and is also part of the EDCTP2 programme supported by the European Union.

## 7 Supplementary Information

### 7.1 Supplementary methods

#### 7.1.1 Mosquito survival (A(t))

To estimate the difference in mosquito survival due to infection status, *E*, we model the mosquito survival data (see Table 1) using the Cox proportional hazards model as follows. The Kaplan-Meier survival curves of the mosquito survival data (Figure S2) show that the proportion of mosquitoes alive does not decline exponentially indicating that the mosquito mortality rate is not constant. Indeed, a previous analysis of a subset of the data found the best fitting survival distribution to be the Gompertz, in which the mosquito mortality rate increases with age (i.e. there is senescence) (14). The baseline hazard, *h*_0_(*t*), was modelled as Gompertzian:

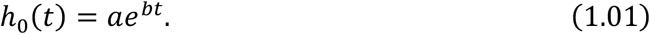

The hazard, *h*_0_(*t*), is modelled accounting for differences in mosquito infection status, *E* (= 1, if infected; = 0, if uninfected), and calculated as,

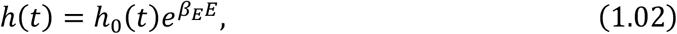

where *β_E_* is a constant. The probability of being alive at time *t*, *A*(*t*), is calculated by integrating the hazard to time t, as follows,

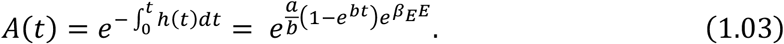

As mosquitoes were removed for dissection or still alive at the end of the study some observations were right censored, that is we do not know the time when death would have occurred. For these observations, the likelihood is given by, *A*(*t*). The probability density function, *f*(*t*), describing the event that death occurs at time *t*, for uncensored observations, is given by,

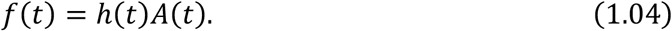

We suppose the infectious blood-fed group is a mix of infected and uninfected mosquitoes. The true infection status of these mosquitoes was not known. The probability of an observable infection in this group is given by *δ*, but we cannot distinguish between mosquitoes that never received any parasites and those that cleared the infection. There may be a potential cost of immunity and tissue damage caused by the ookinete life stage, but given that the impact of malaria infection is still unclear (26), we assume that if the mosquito clears the malaria infection prior to the oocyst life stage, there is no additional mortality hazard due to infection. We account for uncertainty in the true infection status of mosquitoes by marginalising infection status out of the joint distribution as follows,

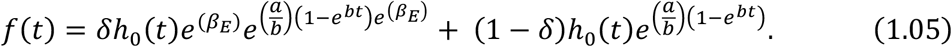

#### 7.1.2 Model fitting

**Figure S1.**
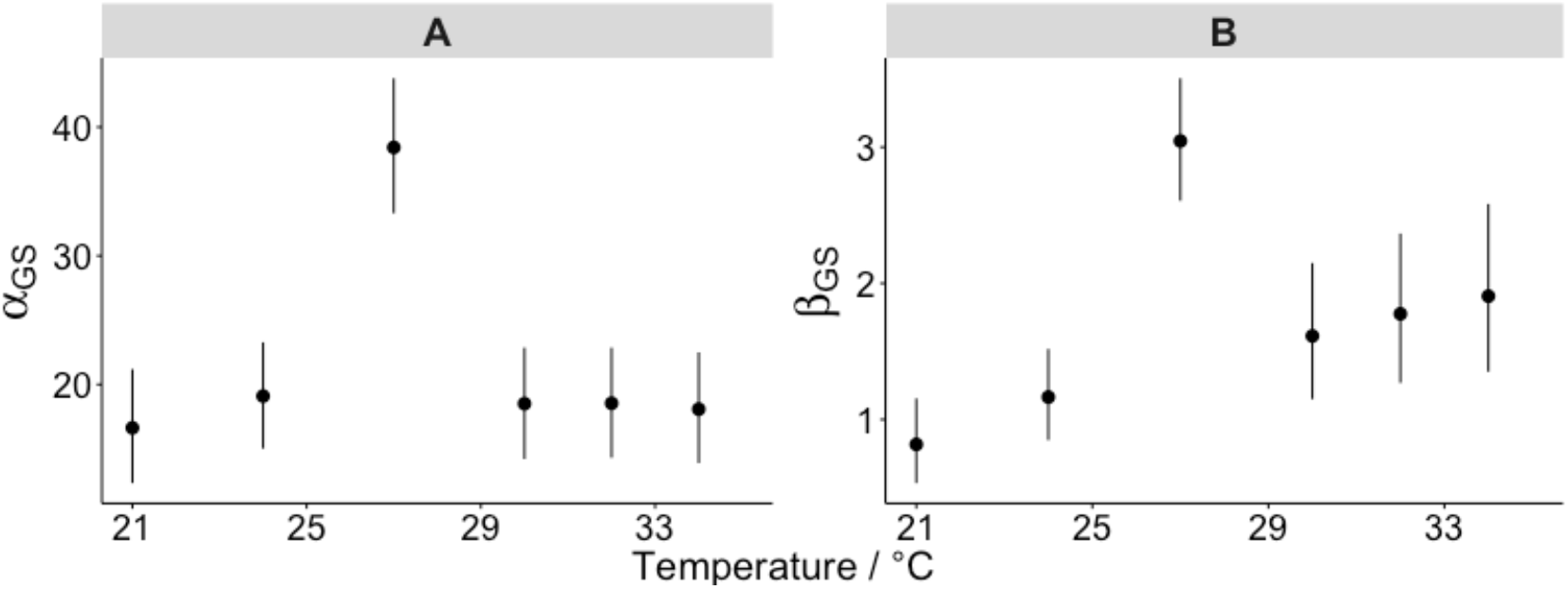
Single temperature model estimates of parasite transition parameters. The gamma distribution posterior values of the development time between inoculation and observed sporozoites for the single temperature models. The MCMC median, 5^th^ and 95^th^ posterior quantiles are represented by the points and vertical lines respectively.

**Figure S2.**
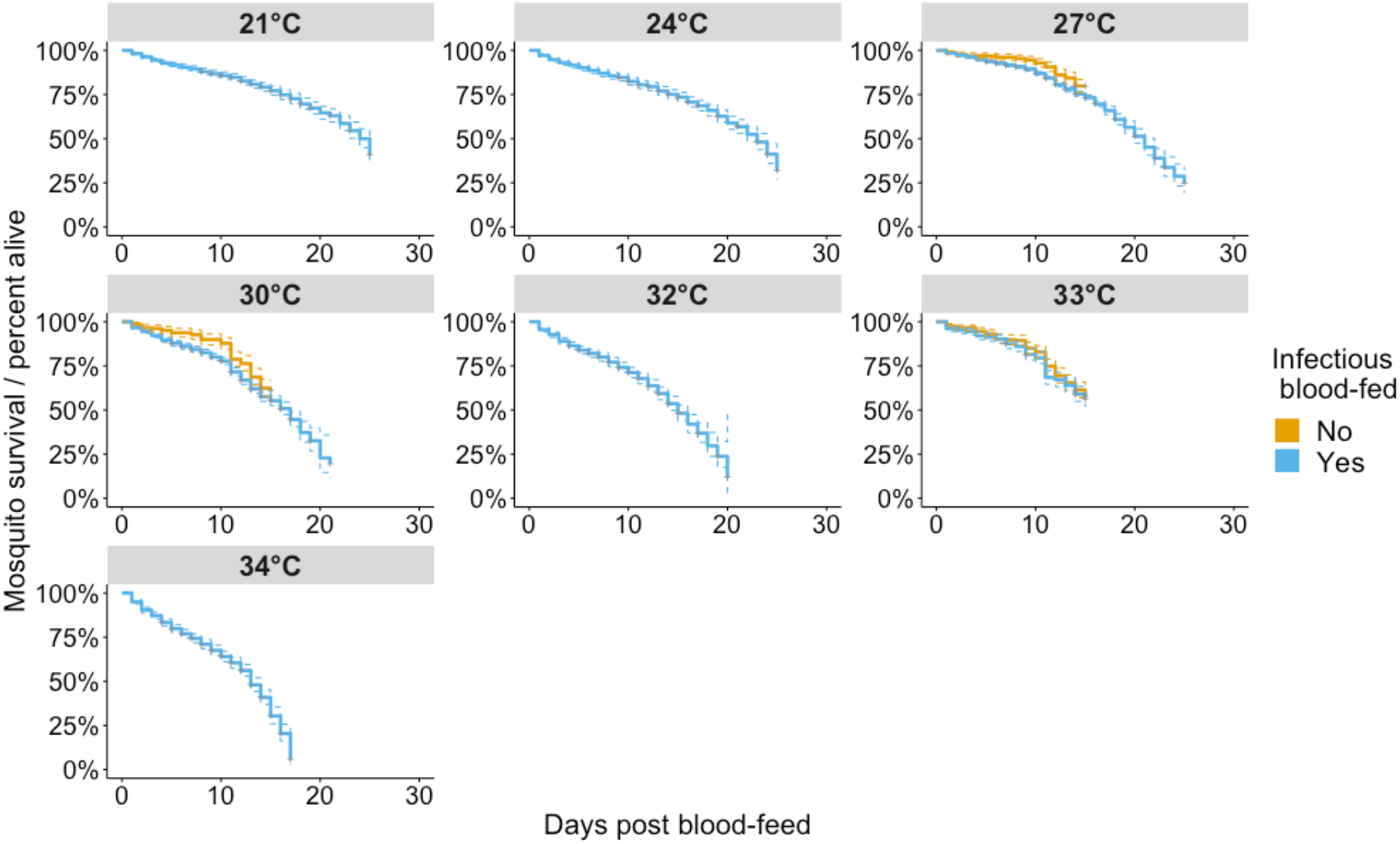
Kaplan Meier mosquito survival curves used in the model fitting.

**Table S1.** Model priors.

| Parameter | Description | Prior parameters<br>(subject to limits in<br>next column) | Parameter<br>limits | Justification of<br>parameterisation<br>(reference) |
| --- | --- | --- | --- | --- |
| **Temperature-independent parameters** |  |  |  |  |
| $\alpha_{OS}$ | Shape parameter for the gamma distribution governing oocyst to sporozoite development time | $\alpha_{OS} \sim N(15.0, 2.5)$ | Lower: 0 | Yields a mean sporozoite development time of 8 days |
| $\beta_{OS}$ | Rate parameter for the gamma distribution governing oocyst to sporozoite development time | $\beta_{OS} \sim N(1.875, 2.5)$ | | |
| $\mu$ | Mean number of G-stage parasites inoculating an infected mosquito | $\mu \sim N(3.0, 2.5)$ | Lower: 0<br>Upper: 50 | The median and mean number of oocysts per mosquito has been found to be approximately 3 (68–71) |
| $k$ | Overdispersion in the number of G-stage parasites inoculating an infected mosquito | $k \sim N(0.1, 2.5)$ | Lower: 0<br>Upper: 20 | The overdispersion of oocysts has been found to be below slightly below 0.1 (70,71) |
| $a$ | Gompertz distribution parameter governing the baseline hazard | $a \sim N(0.05, 2.5)$ | Lower: 0<br>Upper: 5 | Assuming mosquito survival is solely determined by the baseline hazard, the mean prior values result in a median survival time of 10.3 days (0.5 – 30.9 days 95% survival quantiles) |
| $b$ | Gompertz distribution parameter governing the baseline hazard | $b \sim N(0.05, 2.5)$ | | |
| $\beta_E$ | Cox proportional hazards model coefficient of infection status | $\beta_E \sim N(0.5, 1.0)$ | Lower: -5<br>Upper: 5 | The hazard odds ratio of infection is 1.65, given the mean prior value |
| $\sigma_\delta$ | Standard deviation of the delta error term | $\sigma_\delta \sim N(0, 0.25)$ | Lower: 0 | There are no differences between experiments, given the mean prior values |
| $\sigma_{survival}$ | Standard deviation of the Cox model error term | $\sigma_{survival} \sim N(0, 0.25)$ | | |
| **Temperature-dependent parameters** |  |  |  |  |
| $a_{GO,}$ | Shape parameter for the gamma distribution governing G-stage parasite to oocyst development time | $\alpha_{IO} \sim N(15.0, 2.5)$ | Lower: 0 | At the prior mean value of $m_b(0)$ temperature has no effect on the oocyst development rate. The shape and rate priors give a mean oocyst development time of 5 days |
| $m_\beta$ | Temperature-coefficient in eq (2.12) | $m_\beta \sim N(0, 2.5)$ | $m_b$ has no limits | |
| $c_\beta$ | Intercept in eq (2.12) | $c_\beta \sim N(3.0, 2.5)$ | Lower: 0 | |
| $m_\delta$ | Temperature-coefficient in eq (2.13) | $m_\delta \sim N(0, 2.5)$ | | At the prior mean values of temperature has no effect on the transmission probability. Overall these result in a prior mean transmission probability of 0.69 |
| $c_\delta$ | Intercept in eq (2.13) | $c_\delta \sim N(0.8, 2.5)$ | | |
| $\beta_c$ | Temperature coefficient in Cox proportional hazards model (eq (2.15)) | $\beta_c \sim N(0.5, 1.0)$ | Lower: -5<br>Upper: 5 | Mosquitoes have a hazard odds ratio of 1.15 for each 1°C increase in temperature |

**Figure S3.**
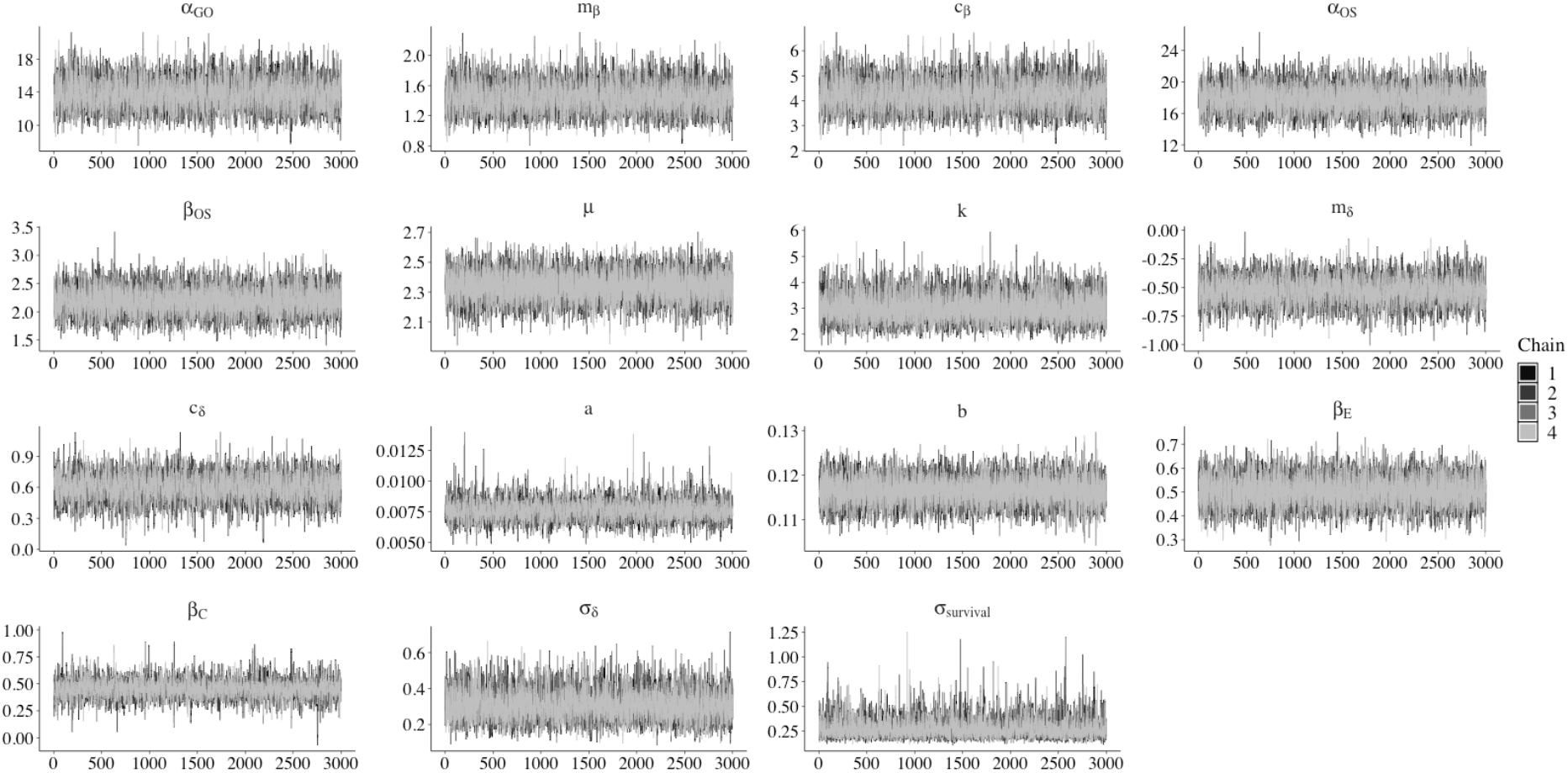
Trace plots of the MCMC parameter sampling for the all temperature model. The horizontal and vertical axis give the iteration and parameter values respectively. Iterations during the warmup are not included.

**Figure S4.**
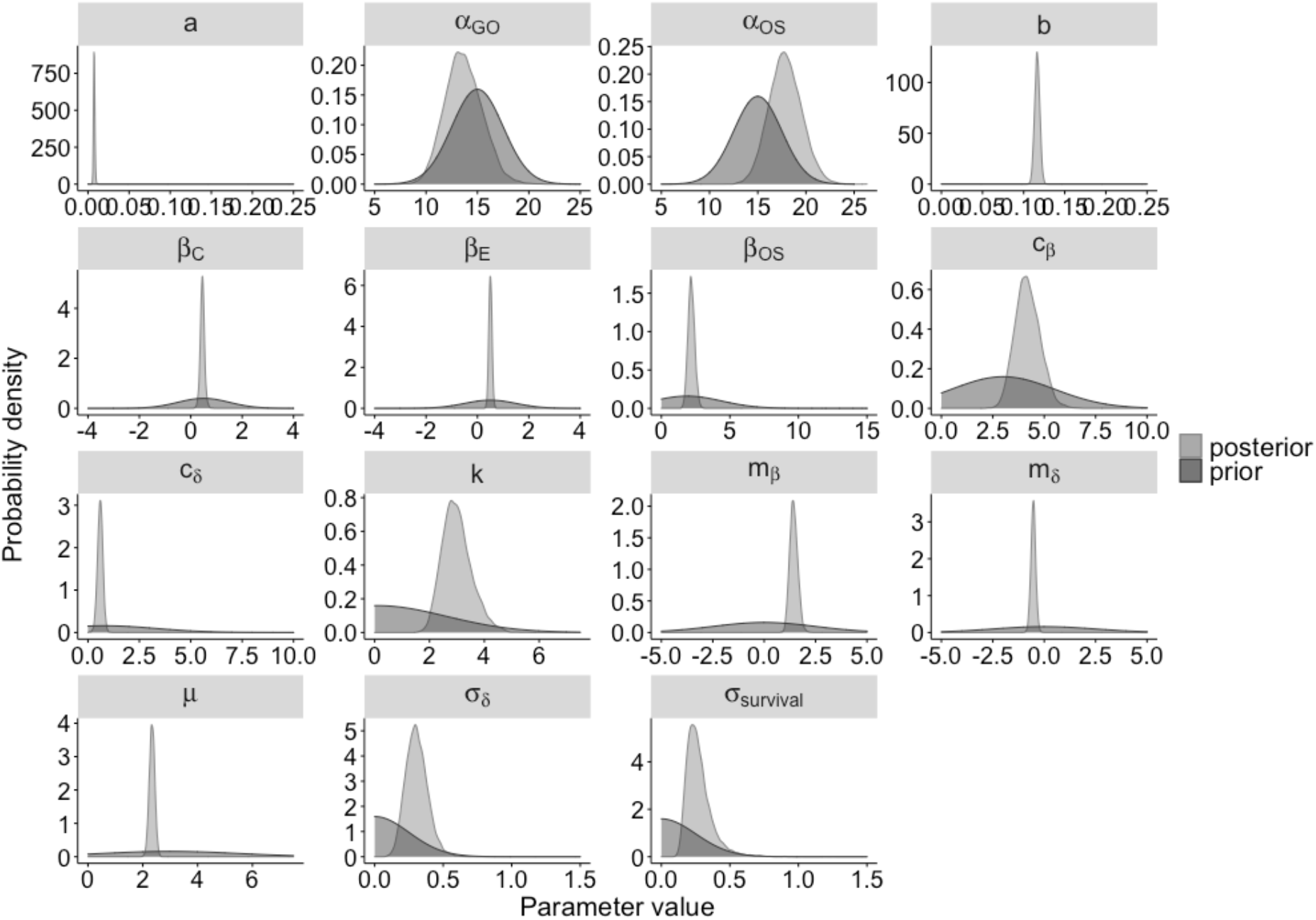
Posterior and prior probability densities. The posterior distributions are estimated by kernel density estimation with a gaussian smoothing kernel.

**Table S2.** All temperature model posterior values.

| Parameter | Mean value | SE of the mean | Standard deviation | 2.5% quantile | 25% quantile | 50% quantile | 75% quantile | 97.5% quantile | ESS | $\hat{R}$ |
| --- | --- | --- | --- | --- | --- | --- | --- | --- | --- | --- |
| $\alpha_{GO}$ | 13.67 | 0.02 | 1.8 | 10.36 | 12.42 | 13.59 | 14.84 | 17.3 | 5808 | 1 |
| $m_\beta$ | 1.44 | 0 | 0.19 | 1.09 | 1.31 | 1.43 | 1.57 | 1.83 | 6182 | 1 |
| $c_\beta$ | 4.19 | 0.01 | 0.6 | 3.1 | 3.78 | 4.16 | 4.59 | 5.42 | 5847 | 1 |
| $\alpha_{OS}$ | 17.87 | 0.02 | 1.69 | 14.69 | 16.71 | 17.82 | 18.99 | 21.29 | 10275 | 1 |
| $\beta_{OS}$ | 2.19 | 0 | 0.24 | 1.75 | 2.03 | 2.18 | 2.35 | 2.68 | 10101 | 1 |
| $\mu$ | 2.34 | 0 | 0.1 | 2.14 | 2.27 | 2.34 | 2.4 | 2.53 | 11620 | 1 |
| $k$ | 3 | 0 | 0.53 | 2.1 | 2.63 | 2.95 | 3.32 | 4.2 | 14393 | 1 |
| $m_\delta$ | -0.53 | 0 | 0.12 | -0.76 | -0.61 | -0.53 | -0.46 | -0.3 | 6435 | 1 |
| $c_\delta$ | 0.59 | 0 | 0.13 | 0.34 | 0.51 | 0.6 | 0.68 | 0.85 | 4900 | 1 |
| $a$ | 0.01 | 0 | 0 | 0.01 | 0.01 | 0.01 | 0.01 | 0.01 | 4772 | 1 |
| $b$ | 0.12 | 0 | 0 | 0.11 | 0.11 | 0.12 | 0.12 | 0.12 | 16156 | 1 |
| $\beta_E$ | 0.5 | 0 | 0.06 | 0.39 | 0.46 | 0.5 | 0.54 | 0.62 | 10200 | 1 |
| $\beta_C$ | 0.45 | 0 | 0.08 | 0.29 | 0.4 | 0.45 | 0.5 | 0.62 | 4727 | 1 |
| $\sigma_\delta$ | 0.31 | 0 | 0.08 | 0.17 | 0.25 | 0.3 | 0.36 | 0.48 | 7127 | 1 |
| $\sigma_{\text{survival}}$ | 0.27 | 0 | 0.09 | 0.15 | 0.21 | 0.26 | 0.31 | 0.51 | 5358 | 1 |

#### 7.1.3 Logistic growth model

The cumulative sporozoite prevalence at time *t*, *g*(*t*), was estimated under the logistic growth model as,

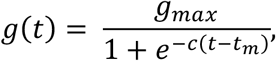

where *g_max_* is the upper asymptote equivalent to the maximum possible sporozoite prevalence (used as an estimate of the human-to-mosquito transmission probability), *t_m_* and *c* are constants that determine the growth rate (27). The binary logistic model was fit to the all dissected mosquitoes (binary coding; 1: sporozoites observed, 0: no sporozoites observed) simultaneously, using non-linear least squares.

### 7.2 Supplementary results

**Figure S5.**
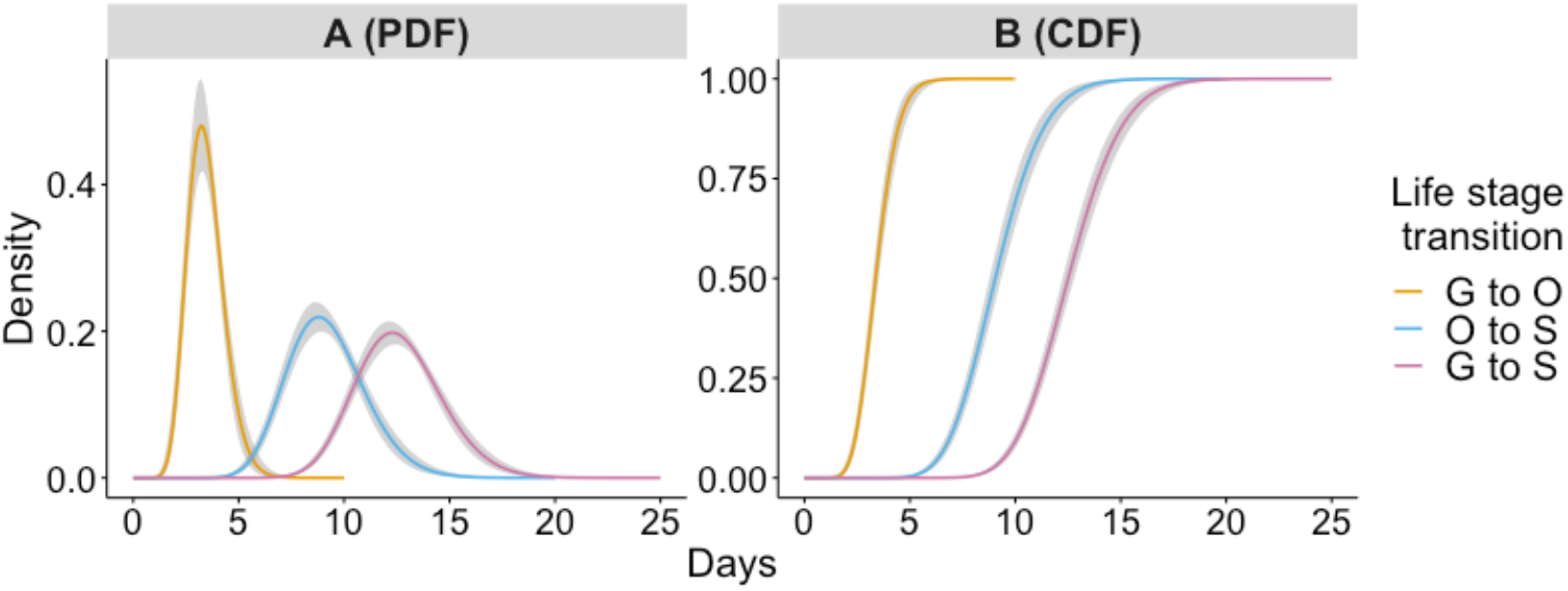
Modelled times required for individual malaria parasites to transition between life stages. Panels A and B show the probability density and the cumulative probability density that an individual malaria parasite within a mosquito (maintained under standard insectary conditions - 27 °C) will transition from a given life stage to the next life stage at a given time post blood-feed. The grey shaded area represents 5%-95% posterior quantiles of estimated distributions.

**Figure S6.**
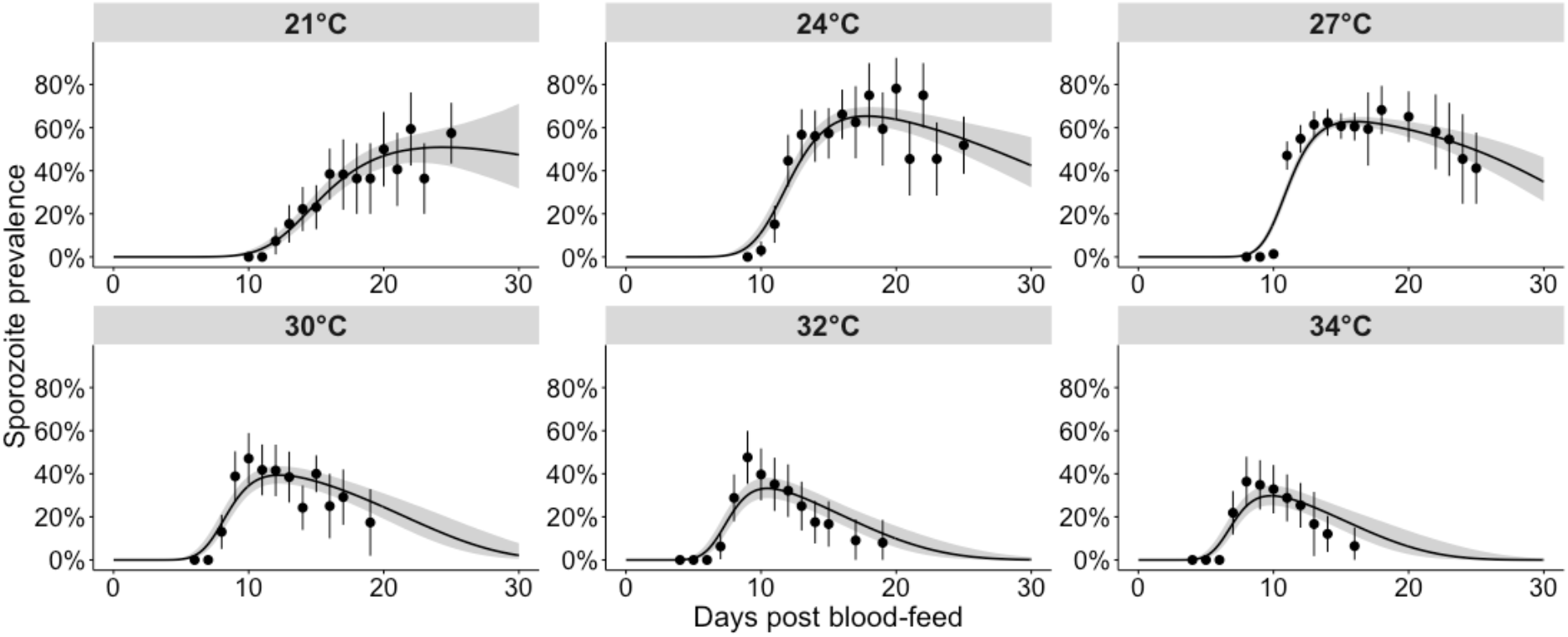
Single temperature model fits of the temporal variation in sporozoite prevalence. Black points: parasite prevalence of the laboratory mosquito data (95% confidence intervals are given by the vertical black lines). The grey shaded area represents the 95% uncertainty intervals of the mean prevalence (posterior predictive means). The black line represents the median of the poster predictive means.

**Figure S7.**
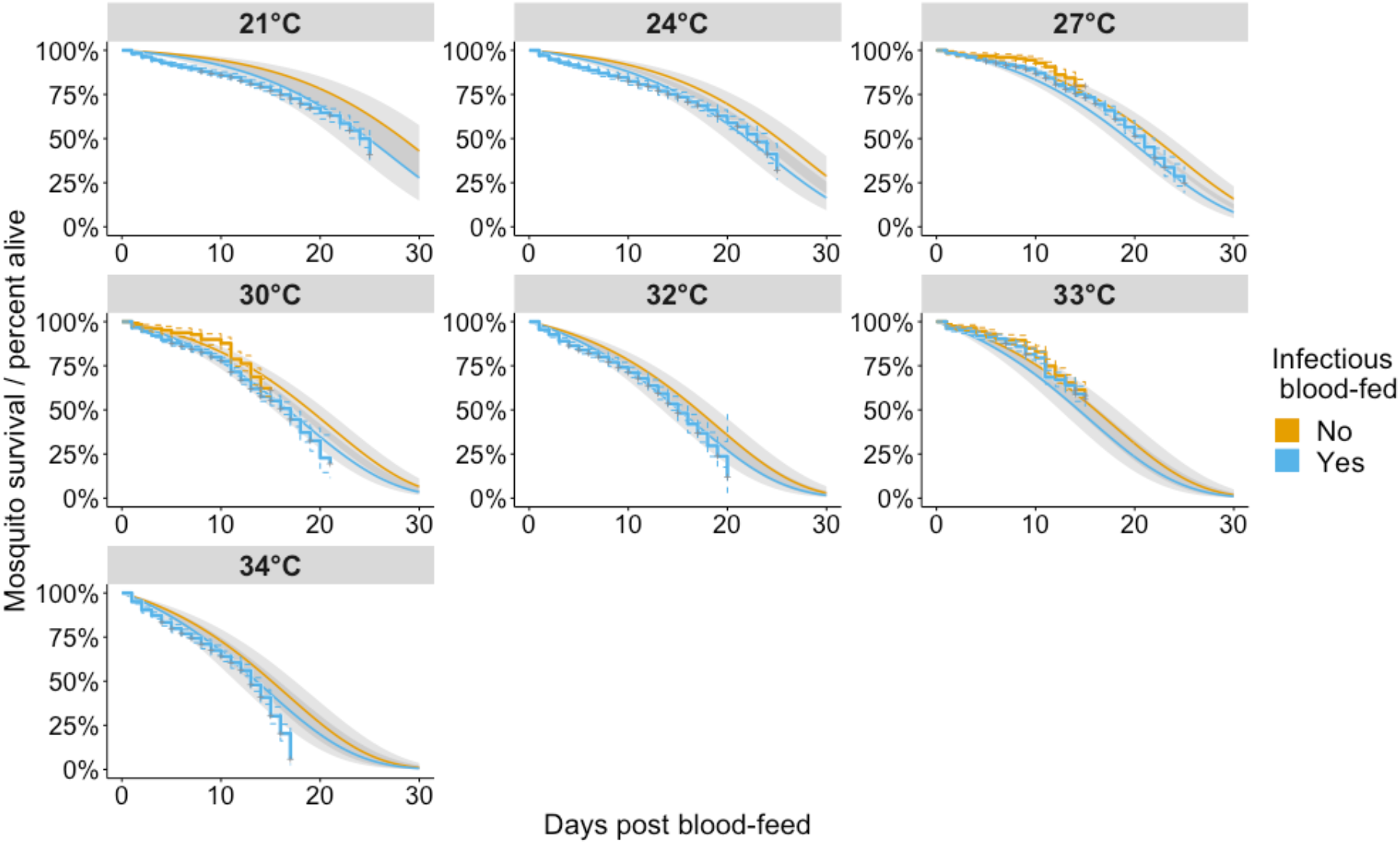
Kaplan-Meier survival curves and all temperature Cox proportional-hazards survival model fit. Kaplan Meier curves are stepped. The shaded area shows the 95% uncertainty intervals of the posterior predictive mean survival probability (A(t)) modelled by the Cox proportional hazards model.

**Figure S8.**
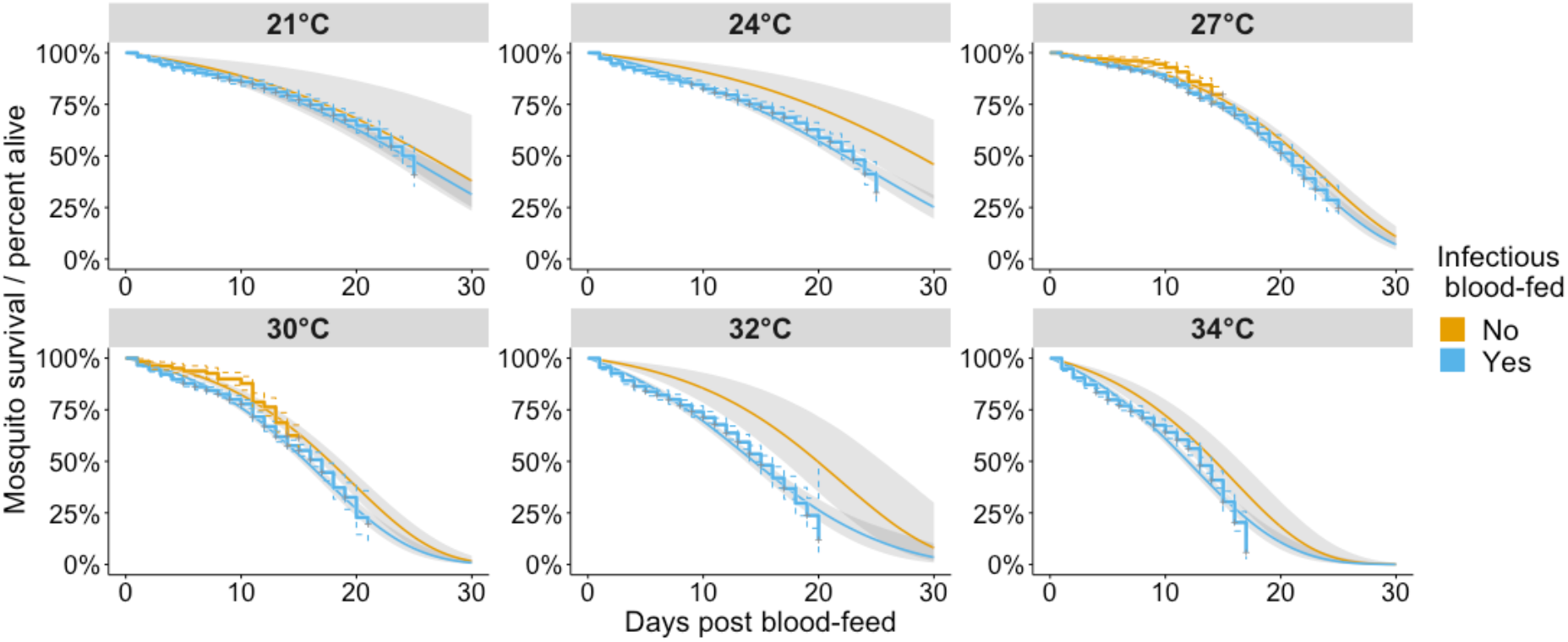
Kaplan Meier survival curves and the single temperature Cox proportional-hazards survival model fits. Kaplan Meier curves are stepped. The shaded area shows the 95% uncertainty intervals of the posterior predictive mean survival probability (A(t)) modelled by the Cox proportional hazards model.

**Figure S9.**
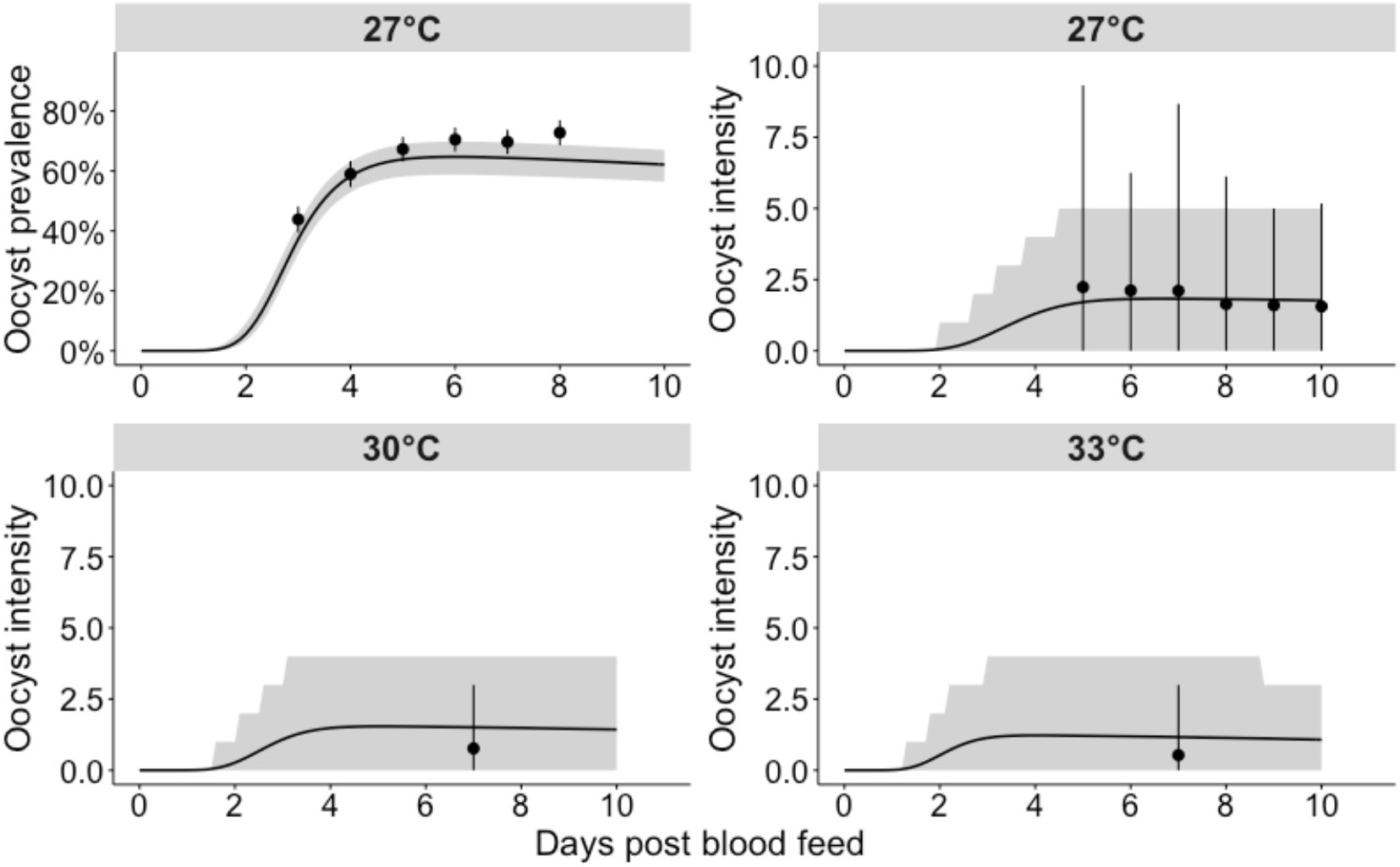
All temperature model fits of the temporal variation in oocyst prevalence and oocyst intensity. Oocyst prevalence plot: black points represent the prevalence of the laboratory mosquito data and the lines represent the 95% confidence intervals.: The grey shaded area represents the 95% uncertainty interval of the posterior predictive mean prevalence. Oocyst intensity plots: black points represent the mean oocyst intensity among all blood-fed mosquitoes and the grey shaded area represents the uncertainty in the oocyst count; the 5% and 95% negative binomial quantiles given the mean posterior predictive mean and mean posterior overdispersion parameter are shown.

**Table S3.** EIP estimates by temperature. The posterior predictive median, 5% and 95% quantiles of the “all temperature model” are given to one decimal place.

| Temperature / °C | EIP<sub>10</sub> | EIP<sub>50</sub> | EIP<sub>90</sub> |
| --- | --- | --- | --- |
| 21 | 12.9 (12.5 - 13.3) | 16.1 (15.5 - 16.8) | 20.4 (19.4 - 21.8) |
| 24 | 9.5 (9.3 - 9.7) | 11.8 (11.6 - 12) | 14.9 (14.5 - 15.2) |
| 27 | 8.2 (8 - 8.3) | 10.2 (10 - 10.4) | 13 (12.7 - 13.3) |
| 30 | 7.5 (7.3 - 7.6) | 9.4 (9.2 - 9.6) | 12 (11.7 - 12.4) |
| 32 | 7.2 (7 - 7.3) | 9 (8.9 - 9.2) | 11.6 (11.3 - 12) |
| 33 | 7 (6.9 - 7.2) | 8.9 (8.7 - 9.1) | 11.5 (11.1 - 11.8) |
| 34 | 6.9 (6.7 - 7.1) | 8.8 (8.6 - 8.9) | 11.3 (11 - 11.7) |

**Figure S10.**
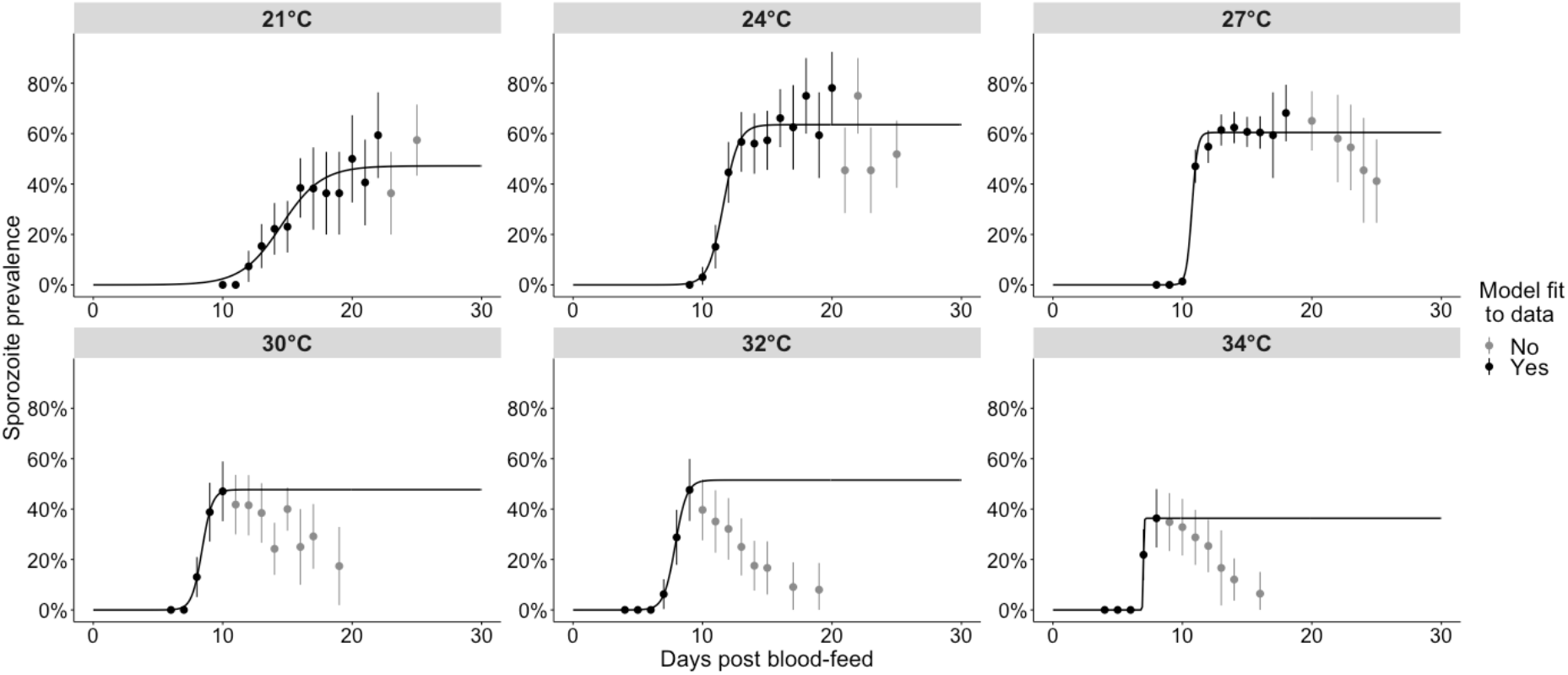
Logistic growth model fits. A logistic growth model (black line) was fit a subset of the sporozoite prevalence data (as determined by the time of peak sporozoite prevalence; black points were included and grey points were excluded).

## References

1. World Health Organization. World malaria report 2019 [Internet]. Geneva; 2019.

2. Bhatt S, Weiss DJ, Cameron E, Bisanzio D, Mappin B, Dalrymple U, et al. The effect of malaria control on Plasmodium falciparum in Africa between 2000 and 2015. Nature. 2015 Sep 16;526:207.

3. Macdonald G. Epidemiological basis of malaria control. Bull World Health Organ. 1956;15(3–5):613.

4. Macdonald G. The analysis of the sporozoite rate. Trop Dis Bull. 1952;49(6).

5. Smith DL, Ellis McKenzie F. Statics and dynamics of malaria infection in Anopheles mosquitoes. Malar J. 2004;3(1):13.

6. Vaughan JA. Population dynamics of Plasmodium sporogony. Trends Parasitol. 2007;23(2):63–70.

7. Pichon G, Awono-Ambene HP, Robert V. High heterogeneity in the number of Plasmodium falciparum gametocytes in the bloodmeal of mosquitoes fed on the same host. Parasitology. 2000;121(2):115–20.

8. Arai M, Billker O, Morris HR, Panico M, Delcroix M, Dixon D, et al. Both mosquito-derived xanthurenic acid and a host blood-derived factor regulate gametogenesis of Plasmodium in the midgut of the mosquito. Mol Biochem Parasitol. 2001;116(1):17–24.

9. Billker O, Lindo V, Panico M, Etienne AE, Paxton T, Dell A, et al. Identification of xanthurenic acid as the putative inducer of malaria development in the mosquito. Nature. 1998;392(6673):289–92.

10. Kuehn A, Pradel G. The coming-out of malaria gametocytes. J Biomed Biotechnol. 2010/01/05. 2010;2010:976827.

11. Clayton AM, Dong Y, Dimopoulos G. The Anopheles innate immune system in the defense against malaria infection. J Innate Immun. 2013/08/28. 2014;6(2):169–81.

12. Gouagna LC, Mulder B, Noubissi E, Tchuinkam T, Boudin C, Boudin C. The early sporogonic cycle of Plasmodium falciparum in laboratory-infected Anopheles gambiae: an estimation of parasite efficacy. Trop Med Int Heal. 1998 Jan 1;3(1):21–8.

13. Vaughan JA, Noden BH, Beier JC. Population dynamics of Plasmodium falciparum sporogony in laboratory-infected Anopheles gambiae. J Parasitol. 1992;78(4):716–24.

14. Shapiro LLM, Whitehead SA, Thomas MB. Quantifying the effects of temperature on mosquito and parasite traits that determine the transmission potential of human malaria. PLOS Biol. 2017 Oct 16;15(10):e2003489.

15. Ohm JR, Baldini F, Barreaux P, Lefevre T, Lynch PA, Suh E, et al. Rethinking the extrinsic incubation period of malaria parasites. Parasit Vectors. 2018;11(1):178.

16. Nikolaev BP. On the influence of temperature on the development of malaria plasmodia in the mosquito. Leningr Pasteur Inst Epidemiol Bacteriol. 1935;2:108–9.

17. Hien DFDS, Dabiré KR, Roche B, Diabaté A, Yerbanga RS, Cohuet A, et al. Plant-Mediated Effects on Mosquito Capacity to Transmit Human Malaria. PLoS Pathog. 2016 Aug 4;12(8):e1005773–e1005773.

18. Shapiro LLM, Murdock CC, Jacobs GR, Thomas RJ, Thomas MB. Larval food quantity affects the capacity of adult mosquitoes to transmit human malaria. Proc R Soc B Biol Sci. 2016 Jul 13;283(1834):20160298.

19. Afrane YA, Little TJ, Lawson BW, Githeko AK, Yan G. Deforestation and vectorial capacity of Anopheles gambiae Giles mosquitoes in malaria transmission, Kenya. Emerg Infect Dis. 2008 Oct;14(10):1533–8.

20. Kærn M, Elston TC, Blake WJ, Collins JJ. Stochasticity in gene expression: from theories to phenotypes. Nat Rev Genet. 2005;6(6):451–64.

21. Paul REL, Bonnet S, Boudin C, Tchuinkam T, Robert V. Aggregation in malaria parasites places limits on mosquito infection rates. Infect Genet Evol. 2007;7(5):577–86.

22. Da DF, Churcher TS, Yerbanga RS, Yaméogo B, Sangaré I, Ouedraogo JB, et al. Experimental study of the relationship between Plasmodium gametocyte density and infection success in mosquitoes; implications for the evaluation of malaria transmission-reducing interventions. Exp Parasitol. 2015;149:74–83.

23. Yassine H, Osta MA. Anopheles gambiae innate immunity. Cell Microbiol. 2010 Jan 1;12(1):1–9.

24. Dong Y, Manfredini F, Dimopoulos G. Implication of the mosquito midgut microbiota in the defense against malaria parasites. PLoS Pathog. 2009/05/08. 2009 May;5(5):e1000423–e1000423.

25. Blanford S, Chan BHK, Jenkins N, Sim D, Turner RJ, Read AF, et al. Fungal Pathogen Reduces Potential for Malaria Transmission. Science (80-). 2005 Jun 10;308(5728):1638 LP – 1641.

26. Ferguson HM, Read AF. Why is the effect of malaria parasites on mosquito survival still unresolved? Trends Parasitol. 2002;18(6):256–61.

27. Paaijmans KP, Blanford S, Chan BHK, Thomas MB. Warmer temperatures reduce the vectorial capacity of malaria mosquitoes. Biol Lett. 2011/12/21. 2012 Jun 23;8(3):465–8.

28. Murdock CC, Paaijmans KP, Bell AS, King JG, Hillyer JF, Read AF, et al. Complex effects of temperature on mosquito immune function. Proceedings Biol Sci. 2012/05/16. 2012 Aug 22;279(1741):3357–66.

29. Pathak AK, Shiau JC, Thomas MB, Murdock CC. Field Relevant Variation in Ambient Temperature Modifies Density-Dependent Establishment of Plasmodium falciparum Gametocytes in Mosquitoes [Internet]. Vol. 10, Frontiers in Microbiology. 2019. p. 2651.

30. Detinova TS, Bertram DS, Organization WH. Age-grouping methods in Diptera of medical importance, with special reference to some vectors of malaria. World Health Organization; 1962.

31. Paaijmans KP, Read AF, Thomas MB. Understanding the link between malaria risk and climate. Proc Natl Acad Sci. 2009 Aug 18;106(33):13844 LP – 13849.

32. Mordecai EA, Paaijmans KP, Johnson LR, Balzer C, Ben-Horin T, Moor E, et al. Optimal temperature for malaria transmission is dramatically lower than previously predicted. Ecol Lett. 2012 Oct 11;16(1):22–30.

33. Teboh-Ewungkem MI, Yuster T. A within-vector mathematical model of Plasmodium falciparum and implications of incomplete fertilization on optimal gametocyte sex ratio. J Theor Biol. 2010;264(2):273–86.

34. Childs LM, Prosper OF. Simulating within-vector generation of the malaria parasite diversity. PLoS One. 2017 May 22;12(5):e0177941–e0177941.

35. Gog JR, Pellis L, Wood JLN, McLean AR, Arinaminpathy N, Lloyd-Smith JO. Seven challenges in modeling pathogen dynamics within-host and across scales. Epidemics. 2015;10:45–8.

36. Murdock CC, Sternberg ED, Thomas MB. Malaria transmission potential could be reduced with current and future climate change. Sci Rep. 2016 Jun 21;6:27771.

37. Bompard A, Da DF, Yerbanga SR, Morlais I, Awono-Ambéné PH, Dabiré RK, et al. High Plasmodium infection intensity in naturally infected malaria vectors in Africa. Int J Parasitol. 2020;

38. Churcher TS, Blagborough AM, Delves M, Ramakrishnan C, Kapulu MC, Williams AR, et al. Measuring the blockade of malaria transmission – An analysis of the Standard Membrane Feeding Assay. Int J Parasitol. 2012;42(11):1037–44.

39. Miura K, Swihart BJ, Deng B, Zhou L, Pham TP, Diouf A, et al. Transmission-blocking activity is determined by transmission-reducing activity and number of control oocysts in Plasmodium falciparum standard membrane-feeding assay. Vaccine. 2016/06/29. 2016 Jul 29;34(35):4145–51.

40. Noden BH, Kent MD, Beier JC. The impact of variations in temperature on early Plasmodium falciparum development in Anopheles stephensi. Parasitology. 2009/04/06. 1995;111(5):539–45.

41. Eling W, Hooghof J, Van De Vegte-Bolmer M, Sauerwein R, Van Gemert GJ. Tropical temperatures can inhibit development of the human malaria parasite Plasmodium falciparum in the mosquito. In: Proceedings of the Section Experimental and Applied Entomology-Netherlands Entomological Society. 2001. p. 151–6.

42. Stewart T, Strijbosch LWG, Moors H, Batenburg P van. A simple approximation to the convolution of gamma distributions. 2007;

43. Stone WJR, Eldering M, van Gemert G-J, Lanke KHW, Grignard L, van de Vegte-Bolmer MG, et al. The relevance and applicability of oocyst prevalence as a read-out for mosquito feeding assays. Sci Rep. 2013 Dec 4;3:3418.

44. David HA, Nagaraja HN. Order statistics. Encycl Stat Sci. 2004;

45. Beier JC. Malaria parasite development in mosquitoes. Annu Rev Entomol. 1998;43(1):519–43.

46. Carpenter B, Gelman A, Hoffman MD, Lee D, Goodrich B, Betancourt M, et al. Stan: A probabilistic programming language. J Stat Softw. 2017;76(1).

47. Hoffman MD, Gelman A. The No-U-Turn sampler: adaptively setting path lengths in Hamiltonian Monte Carlo. J Mach Learn Res. 2014;15(1):1593–623.

48. Shaw WR, Holmdahl IE, Itoe MA, Werling K, Marquette M, Paton DG, et al. Multiple blood feeding in mosquitoes shortens the *Plasmodium falciparum* incubation period and increases malaria transmission potential. bioRxiv. 2020 Jan 1;2020.03.24.991356.

49. Chege GMM, Beier JC. Effect of Plasmodium falciparum on the Survival of Naturally Infected Afrotropical Anopheles (Diptera: Culicidae). J Med Entomol. 1990 Jul 1;27(4):454–8.

50. Brugman VA, Kristan M, Gibbins MP, Angrisano F, Sala KA, Dessens JT, et al. Detection of malaria sporozoites expelled during mosquito sugar feeding. Sci Rep. 2018;8(1):7545.

51. Billingsley PF, Hodivala KJ, Winger LA, Sinden RE. Detection of mature malaria infections in live mosquitoes. Trans R Soc Trop Med Hyg. 1991;85(4):450–3.

52. Snow RW, Gouws E, Omumbo J, Rapuoda B, Craig MH, Tanser FC, et al. Models to predict the intensity of Plasmodium falciparum transmission: applications to the burden of disease in Kenya. Trans R Soc Trop Med Hyg. 1998 Nov 1;92(6):601–6.

53. Gosoniu L, Vounatsou P, Sogoba N, Maire N, Smith T. Mapping malaria risk in West Africa using a Bayesian nonparametric non-stationary model. Comput Stat Data Anal. 2009;53(9):3358–71.

54. Reiner RC, Geary M, Atkinson PM, Smith DL, Gething PW. Seasonality of Plasmodium falciparum transmission: a systematic review. Malar J. 2015;14(1):343.

55. Gething PW, Van Boeckel TP, Smith DL, Guerra CA, Patil AP, Snow RW, et al. Modelling the global constraints of temperature on transmission of Plasmodium falciparum and P. vivax. Parasit Vectors. 2011;4(1):92.

56. Ryan SJ, Lippi CA, Zermoglio F. Shifting transmission risk for malaria in Africa with climate change: a framework for planning and intervention. Malar J. 2020;19(1):170.

57. Suh E, Grossman MK, Waite JL, Dennington NL, Sherrard-Smith E, Churcher TS, et al. The influence of feeding behaviour and temperature on the capacity of mosquitoes to transmit malaria. Nat Ecol Evol. 2020;

58. Paaijmans KP, Blanford S, Bell AS, Blanford JI, Read AF, Thomas MB. Influence of climate on malaria transmission depends on daily temperature variation. Proc Natl Acad Sci. 2010 Aug 24;107(34):15135 LP – 15139.

59. Mordecai EA, Caldwell JM, Grossman MK, Lippi CA, Johnson LR, Neira M, et al. Thermal biology of mosquito-borne disease. Ecol Lett. 2019 Oct 1;22(10):1690–708.

60. Tanser FC, Sharp B, le Sueur D. Potential effect of climate change on malaria transmission in Africa. Lancet. 2003;362(9398):1792–8.

61. Siraj AS, Santos-Vega M, Bouma MJ, Yadeta D, Carrascal DR, Pascual M. Altitudinal Changes in Malaria Incidence in Highlands of Ethiopia and Colombia. Science (80-). 2014 Mar 7;343(6175):1154 LP – 1158.

62. Franklinos LH V, Jones KE, Redding DW, Abubakar I. The effect of global change on mosquito-borne disease. Lancet Infect Dis. 2019;19(9):e302–12.

63. Gething PW, Smith DL, Patil AP, Tatem AJ, Snow RW, Hay SI. Climate change and the global malaria recession. Nature. 2010;465(7296):342–5.

64. Sinden RE, Dawes EJ, Alavi Y, Waldock J, Finney O, Mendoza J, et al. Progression of Plasmodium berghei through Anopheles stephensi is density-dependent. PLoS Pathog. 2007 Dec 28;3(12):e195–e195.

65. Mendes AM, Awono-Ambene PH, Nsango SE, Cohuet A, Fontenille D, Kafatos FC, et al. Infection intensity-dependent responses of Anopheles gambiae to the African malaria parasite Plasmodium falciparum. Infect Immun. 2011;79(11):4708–15.

66. Lyimo EO, Koella JC. Relationship between body size of adult Anopheles gambiae sl and infection with the malaria parasite Plasmodium falciparum. Parasitology. 1992;104(2):233–7.

67. Habtewold T, Sharma AA, Wyer CAS, Masters EKG, Windbichler N, Christophides GK. Plasmodium oocysts respond with dormancy to crowding and nutritional stress. bioRxiv. 2020 Jan 1;2020.03.07.981951.

68. Pringle G. A quantitative study of naturally-acquired malaria infections in Anopheles gambiae and Anopheles funestus in a highly malarious area of East Africa. Trans R Soc Trop Med Hyg. 1966;60(5):626–32.

69. Rosenberg R, Andre RG, Somchit L. Highly efficient dry season transmission of malaria in Thailand. Trans R Soc Trop Med Hyg. 1990 Jan 1;84(1):22–8.

70. Taylor LH. Infection rates in, and the number of Plasmodium falciparum genotypes carried by Anopheles mosquitoes in Tanzania. Ann Trop Med Parasitol. 1999 Sep 1;93(6):659–62.

71. Billingsley PF, Medley GF, Charlwood D, Sinden RE. Relationship between prevalence and intensity of Plasmodium falciparum infection in natural populations of Anopheles mosquitoes. Am J Trop Med Hyg. 1994;51(3):260–70.

